# Combination of in vitro and ex vivo infection models highlights the role of NAD-dependent ADP-ribosylation in Influenza A virus infection

**DOI:** 10.1101/2025.10.14.682270

**Authors:** Florentine Jacolin, Anne Aublin-Gex, Manon Vellet, Didier Décimo, Maxime Lepetit, Titouan Denecheau, Lola Canus, Eva Ogire, Adeline Cezard, Clémence Jacquemin, Garance Potelet, Severine Croze, Joel Lachuer, Vincent Lotteau, Olivier Diaz, Guillaume Marcy, Pierre-Olivier Vidalain, Mustapha Si-Tahar, Cyrille Mathieu, Laure Perrin-Cocon

## Abstract

Influenza A viruses (IAV) remain a major public health concern, causing an estimated 650 000 deaths annually. Although vaccines are available, vaccination coverage is limited and must be repeated annually due to the genetic variability of these viruses. Furthermore, although direct acting antivirals are available, they must be administered as early as possible after infection to be effective. In this context, understanding the metabolic alterations associated with the replication and pathogenesis of IAV could contribute to the development of treatments that promote virus clearance and patient recovery. In this study, we identified the intracellular level of nicotinamide adenine dinucleotide (NAD) as a key factor in the replication of IAV. The depletion of NAD using nicotinamide phosphoribosyltransferase (NAMPT) inhibitors or the administration of the antimetabolite 6-aminonicotinamide (6-AN) suppressed IAV replication both *in vitro* and *ex vivo*. Using a model of infected murine organotypic lung cultures (mOLCs) combined with untargeted metabolomic profiling and spatial transcriptomic analyses, our research also revealed that IAV-infection decreased NAD^+^ levels in mOLCs, while PARPs, a group of NAD^+^-consuming enzymes, were upregulated. Pharmacological inhibition of the mono-ADP-ribosyl transferase (mono-ART) activity of PARPs restricted IAV infection *ex vivo*, suggesting that NAD^+^ sustains the proviral activity of these enzymes. Overall, our study identifies NAD metabolism as a central regulator of IAV infection and highlights NAMPT and PARP mono-ART activities as promising antiviral targets.

**Significance Statement:** Viruses are intracellular parasites that rely on the host cellular metabolism for their replication. Our results demonstrate that Influenza A virus (IAV) replication depends on NAD availability, an enzyme cofactor essential for redox metabolic reactions and a substrate for NAD^+^-consuming enzymes. Using a novel *ex vivo* infection model of adult mouse lung tissue, we found that IAV infection leads to increased expression of NAD^+^-consuming PARP enzymes and NAD^+^ depletion. Inhibitors of NAD biosynthesis and of the mono-ADP-ribosyl transferase activity of PARPs restrict viral replication and the production of infectious particles. These findings highlight a proviral role for PARP activity and identify NAMPT and PARP inhibitors as potential host-directed antivirals against IAV.

## Introduction

Influenza A viruses (IAVs) are responsible for seasonal epidemics of respiratory infections in humans, which along with Influenza B, result in approximately one billion cases worldwide each year (1). In some cases, the disease progresses into severe pneumonia and multi-organ failure (2). Despite the availability of vaccines and antiviral treatments, seasonal epidemics lead to an estimated 4 million severe cases and up to 650 000 deaths annually (1). Approved antivirals against IAV target different viral proteins: M2, which forms a proton channel essential for viral entry (Amantadine, Rimantadine), the neuraminidase, which enables viral shedding (Oseltamivir, Zanamivir), and the endonuclease of the viral RNA-dependent RNA polymerase (RdRp) (Baloxavir marboxil) (3). However, the effectiveness of these drugs in reducing symptoms, disease severity, disease and hospitalization duration is limited if they are not administered rapidly after infection, and generally within 48 hours (4). Moreover, the emergence of resistance variants has been detected in 5 to 10% of the patients treated with Baloxavir, and circulating strains are now resistant to Amantadine and Rimantadine. Thus, IAVs remain a public health issue due to their ability to mutate and reassort, leading to the emergence of variants that may escape existing vaccines and antiviral drugs.

Like other viruses, IAVs depend on host cell resources to replicate and disseminate, hijacking cellular metabolism. A deep understanding of IAV-induced metabolic dysregulations could lead to the development of new therapeutic strategies. IAV infection especially enhances glycolytic activity in epithelial cells and mouse lungs (5–8). This glycolytic shift is notably driven by HIF-1α activation which induces glycolytic enzymes, especially hexokinase 2, the first rate-limiting enzyme of glycolysis, and promotes H1N1 viral replication and the host inflammatory response *in vitro* (8–11). Conversely, inhibiting glycolysis with 2-deoxy-glucose (2-DG) in A549 cells reduced viral progeny in a dose-dependent manner (8, 12), and 2-DG administration in vivo reduced viral mRNA expression and lung lesions in IAV H1N1 infected mice (10). These results were further supported by a recent study showing that treatment with either 2-DG or the HIF-1α inhibitor PTX-478 reduced lung viral load, lactate production, and lung damage in IAV-infected mice (11). The metabolic changes observed during infection may arise from viral replication itself or from the host response to infection. For instance, danger signals released by infected cells, such as HMGB1, can engage Toll-like receptor 4 (TLR4), thereby activating the innate immune response and driving metabolic reprogramming of immune cells (13). Consistently, metabolite levels fluctuate in mice following non-lethal IAV challenge, with variations reflecting both the stage of infection and the host response (6, 7). Harnessing these metabolic alterations could therefore not only inhibit viral replication but also improve the management of acute inflammation in severe cases. Accordingly, previous work showed that treatment with TCA cycle metabolites succinate or cis-aconitate, inhibits IAV infection *in vitro*, *ex vivo* and *in vivo*, reducing viral growth, inflammation, and immune cell activation, significantly improving survival after a lethal challenge of IAV infection (14, 15).

To identify new enzymatic pathways crucial for IAV life cycle, we screened a chemical library of metabolic modulators in a respiratory cell line. Drugs targeting the salvage pathway of nicotinamide adenine dinucleotide (NAD), a key redox cofactor involved in many metabolic pathways, such as glycolysis, exerted potent antiviral effects in both *in vitro* and *ex vivo* models of IAV infection. Further investigation of the interplay between NAD metabolism and IAV infection highlighted the role of poly-ADP ribosyl transferases (PARPs), a family of NAD-consuming enzymes. Overall, our results demonstrate that IAV replication critically depends on NAD availability at different stages of the infection.

## Results

### Screening a library of metabolic regulators reveals IAV’s dependence on NAD biosynthesis for replication

To identify metabolic pathways involved in IAV replication, the 493 compounds of a metabolism-related chemical library were screened for their antiviral activity. A549 cells were pretreated for 24 h with 10 µM of each compound, or DMSO alone as a control, before being infected with IAV H3N2 Scotland strain (Fig. 1A). Neuraminidase activity (NA) was measured at 48 h post-infection (hpi) in cell supernatants as a proxy of viral production. Cell viability was measured by counting nuclei after Hoechst staining of adherent cells at 48 hpi. Compounds that reduced the cell number by more than 20% were considered as toxic and were excluded (16% of all tested compounds; Fig. 1B). Baloxavir, which was used as a reference antiviral in the screen, restrained the secretion of NA by more than 90% compared to DMSO-treated cells (*SI appendix*, Fig. S1A-B). To evaluate the robustness of the assay, the Z’ coefficient was calculated as described by Zhang et al., using the NA activity signal from Baloxavir-vs DMSO-treated cells (16). The Z’ coefficient was close to 0.5, indicating that the assay is reliable, although selected hits require secondary validation (*SI appendix*, Fig. S1A-B). Metabolic regulators were considered as potential IAV inhibitors when the released NA activity was reduced by at least 60% compared to DMSO-treated cells, which is outside the bandwidth of positive controls (mean ± 3SD; i.e. 99.73% confidence limit; *SI appendix*, Fig. S1A). Ten compounds filled these criteria (*SI appendix*, Fig. S1C) and two of these hits, FK-866 (APO866) and STF-118804 (Fig. 1B), are inhibitors of nicotinamide phosphoribosyltransferase (NAMPT), a key enzyme in NAD biosynthesis. NAMPT is a rate-limiting enzyme converting nicotinamide (NAM) into nicotinamide mononucleotide (NMN), which is the main precursor of NAD (Fig. 1C). Starting from 10 µM, deescalating doses were tested and confirmed that these two NAMPT inhibitors (NAMPTi) inhibited the production of NA in a dose-dependent manner in IAV-infected A549 cells. Dose-response curves centered on the EC50 were repeated and the upper concentration retaining full antiviral activity for each compound was considered as optimal. (*SI appendix*, Fig. S1D-E). At optimal concentrations of 80 nM for FK-866 and 2 µM for STF-118804, these drugs strongly reduced intracellular NAD levels after 24 h of treatment (Fig. 1D) without affecting the cell count (*SI appendix*, Fig. S1F) at the time when cells were infected with IAV. The impact of NAMPTi on IAV replication was then analyzed by quantifying the viral M segment by RT-qPCR at 48 hpi. Pretreatment with STF-118804 (2 µM) or FK-866 (80 nM) led to a 60–75% reduction in M-segment copy number (Fig. 1E) and resulted in a potent inhibition (around 80%) of the production of viral infectious particles (Fig. 1F). Quantification of total RNA content in infected A549 cells confirmed the absence of significant toxicity of FK-866 and STF-118804 at these concentrations (*SI appendix*, Fig. S1G). Finally, we showed that pretreatments with these drugs impaired early steps of viral infection as they reduced by more than 50% the mean expression of viral nucleoprotein (NP) and non-structural protein 1 (NS1) at 4 hpi (*SI appendix*, Fig. S1H-I). In comparison, pretreatment with Baloxavir which inhibits cap-snatching for viral transcripts, reduced NP expression by approximately 90% and abrogated NS1 expression at the same early time point.

**Figure 1.**
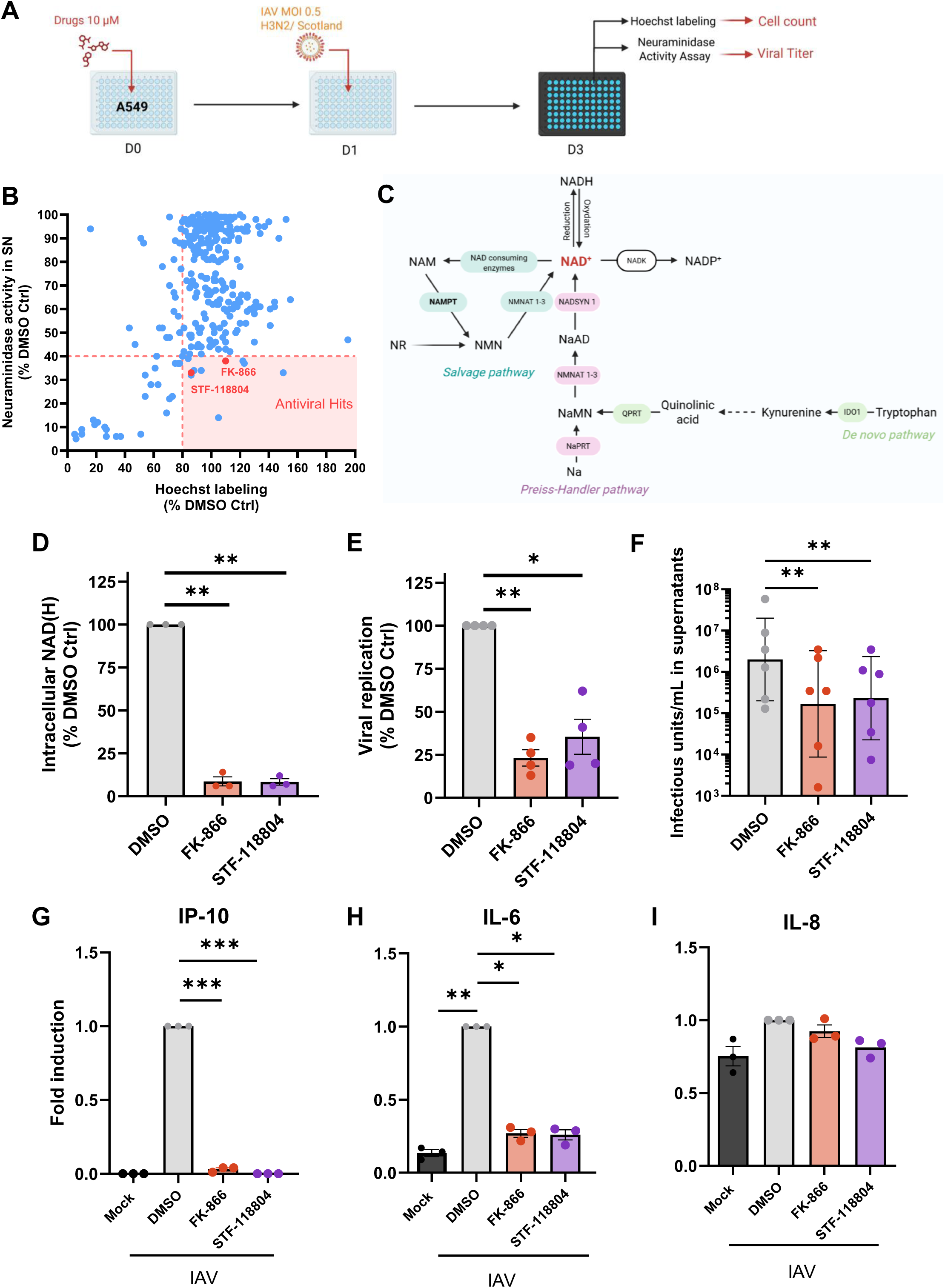
Inhibitors of NAD biosynthesis restrain IAV infection, production and inflammatory antiviral response. (A) Screening strategy of a library of 493 metabolic regulators: A549 cells were seeded and pretreated for 24 h with 10 µM of each compound. Influenza A H3N2/Scotland virus was introduced at MOI 0.5 without medium change, and 48 h post-infection viral production and cell number were respectively determined by neuraminidase (NA) activity assay in supernatants and Hoechst labeling. (B) NA activity and Hoechst labeling were compared to infected untreated control (DMSO Ctrl) and plotted for the 493 compounds. (C) NAD biosynthesis pathway. Most of NAD^+^ is recycled through the salvage pathway using nicotinamide (NAM), nicotinamide riboside (NR) or nicotinamide mononucleotide (NMN) as precursors. In the salvage pathway, the nicotinamide phosphoribosyltransferase (NAMPT) is the rate limiting enzyme. NAD^+^ can also be synthesized from nicotinic acid (Na) through the Preiss-Handler Pathway or in some organs, from tryptophan by the *de novo* pathway. (D-I) A549 cells were pretreated for 24 h with NAMPT inhibitors at 80 nM for FK-866 or 2 µM for STF-118804 or control DMSO solvent, before being infected with IAV MOI 0.5 for 48 h. Intracellular NAD(H) content was quantified at the end of the pretreatment (D). RNA expression level of M viral segment was quantified at 48 hpi by RT-qPCR and normalized with mRNA quantities of Rplp0 cellular gene (E). The production of infectious viral particles in cells supernatants was determined at 48 hpi by TCID_50_ assays on MDCK cells (F). Cytokines secretion was quantified in A549 supernatants collected at 48 hpi by cytometric bead array against IP-10 (G), IL-6 (H) and IL-8 (I) proteins. Data represent means ± SEM or geometrical means ± SD for log_10_ data of at least 3 independent experiments.

IAV infection triggers a strong inflammatory response that contributes to viral pathogenesis and, when uncontrolled, to a cytokine storm that can lead to organ dysfunction and tissue damage (17). Interestingly, NAMPTi prevented the induction by IAV of the interferon-induced protein 10 (IP-10) and strongly reduced IL-6 induction in infected A549 cells (Fig. 1G-H). IL-8 secretion levels were not affected neither by the infection nor by NAMPTi treatment (Fig. 1I). In conclusion, two structurally distinct NAMPTi showed an antiviral activity characterized by the inhibition of IAV replication, infectious viral particles production and IAV-induced antiviral cytokines.

To validate the effect of NAMPTi on host cell metabolism, we analyzed the glycolytic activity and respiration of cells treated or not with FK-866 using an extracellular flux analyzer (XF analyzer). As a control, cells were treated in parallel with the antimetabolite 6-aminonicotinamide (6-AN), which generates non-reducible NAD/NADP analogs, thereby decreasing the pool of NAD (*SI appendix*, Fig. S2A) and interfering with enzymatic reactions dependent on this cofactor, such as glycolysis. Measurement of the proton efflux rate (PER) associated with lactate secretion revealed strong inhibition of the glycolytic rate at 24 and 48 h of treatment with 6-AN, and after 48 h with FK-866 (Fig. 2A-B). Compensatory glycolysis, which corresponds to the ability of cells to increase glycolysis in response to the inhibition of respiration, was markedly more reduced than basal glycolysis (Fig. 2C-D). This is a sign of reduced metabolic flexibility in cells under conditions of high energy demand. The addition of NAM in excess completely abrogated the metabolic effects of 6-AN, confirming that the supply of a NAD precursor restores glycolytic activity. Interestingly, only FK-866 reduced the basal oxygen consumption rate (OCR), suggesting significantly different effects of 6-AN and FK-866 on mitochondrial respiration (Fig. 2E-F). Overall, our results showed that FK-866 has a strong effect on NAD-dependent enzymatic pathways involved in central carbon metabolism, which includes the glycolysis, tricarboxylic acid (TCA) cycle, and the pentose phosphate pathway.

**Figure 2.**
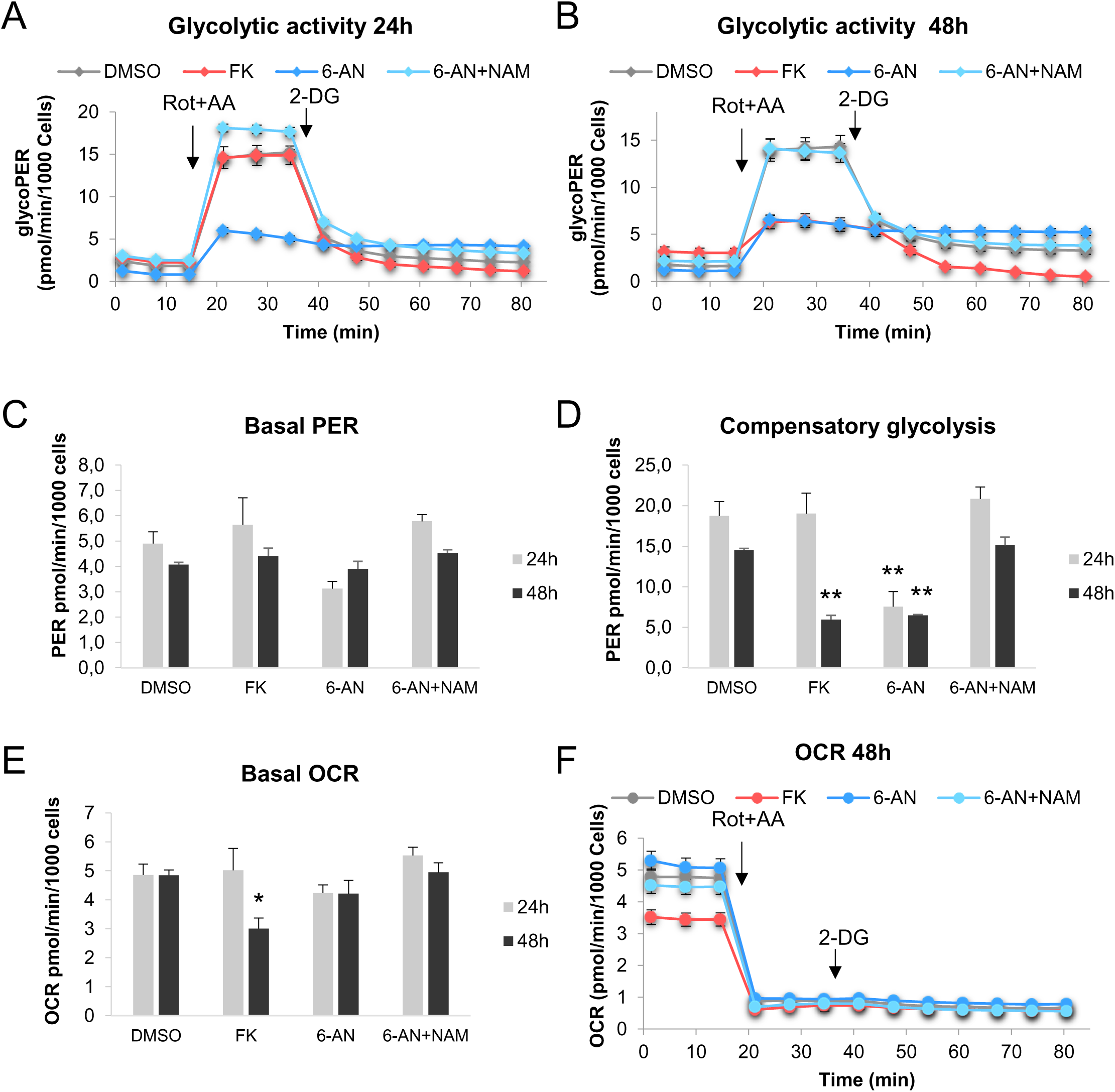
Metabolic activity of A549 cells treated with NAMPTi or 6-AN. Cells were seeded in 24 well-XF plates and treated for 24 or 48h with 80 nM FK-866 (FK), 100 µM 6-AN with or without 500 µM NAM, or control solvent (DMSO). Cells were washed to perform Glycolytic Rate Assay. (A, B) Glycolytic Proton Efflux Rate (PER) profiles of a representative experiment. Mean ± SEM of 5 replicates (C) Basal glycolytic activity. Mean ± SEM basal PER for 3 independent experiments performed in 5 replicates. (D) Compensatory glycolytic activity measured after injection of Rotenone + Antimycin (Rot+AA). Mean ± SEM for 3 independent experiments performed in 5 replicates. (E) Oxygen Consumption Rate (OCR) at basal stage. Mean ± SEM for 3 independent experiments performed in 5 replicates. (F) OCR profile of a representative experiment. Mean ± SEM of 5 replicates.

To further investigate the role of NAD metabolism in IAV replication, we also evaluated the impact of 6-AN treatment following IAV infection in both A549 and BEAS-2B (non-tumoral bronchial) cells. Treatment with 6-AN reduced the NAD pool by 25% in A549 and by 50% in BEAS-2B cells without affecting cell viability. This depletion resulted in a 50% (A549) to 70% (BEAS-2B) reduction in IAV replication and infectious particle production at 24 hpi, an effect that was reversed by the exogenous addition of NAM (*SI appendix*, Fig. S2). These findings are consistent with previous reports describing the broad-spectrum antiviral properties of 6-AN (18–22). Collectively, our results demonstrate that a sufficient amount of NAD is required to support productive IAV infection in A549 and BEAS-2B cells.

### NAD pathway modulators reduce IAV replication and viral production in murine lung tissue

To evaluate the antiviral effect of NAMPTi in a more physiological system, we developed an *ex vivo* model of murine organotypic lung cultures (mOLCs) that supports the infection by IAV. mOLCs were prepared as previously described from adult lung tissue, which was sliced and then cultured at the air-liquid interface to mimic the lung environment *in vivo* (Fig. 3A) (23, 24). Organotypic cultures maintain the structure of tissues, preserve all cell types, especially immune cells, and the intercellular connections within the tissue. The mOLCs were infected with 2 000 or 20 000 PFU of IAV (H3N2 Scotland strain), and viral replication was monitored at 48 hpi by RT-qPCR. Results showed that IAV viral replication in mOLCs was dependent on the initial viral inoculum (Fig. 3B). A kinetic study revealed an initial burst in IAV replication in the first 24 hpi (1.5 log_10_ increase), and maintenance of viral RNA for at least 3 days (Fig. 3C). Infectious foci were detected in infected mOLC by viral NP immunostaining at 48 hpi (*SI appendix*, Fig. S3A). In addition, infectious viral particles were quantified by TCID_50_ assays in mOLC lysates. Viral inoculum was detected at 0 hpi and a 2 log_10_-increase in viral titer was observed at 24 hpi, indicating efficient viral replication *ex vivo* (*SI appendix*, Fig. S3B). MTT tetrazolium metabolization, used as an indicator of tissue viability, showed no difference between infected and uninfected mOLCs, and the signal remained stable for up to 3 days (*SI appendix*, Fig. S3C). Finally, the addition of Baloxavir, a reference antiviral treatment, in the culture medium immediately after infection potently inhibited viral replication in a dose-dependent manner (*SI appendix*, Fig. S3D), with a 3 log_10_ reduction in IAV genome copy number at 100 nM (Fig. 3D). Altogether, these results indicate that mOLC is a relevant model that supports IAV replication and viral production over time and allows the evaluation of antiviral molecules.

**Figure 3.**
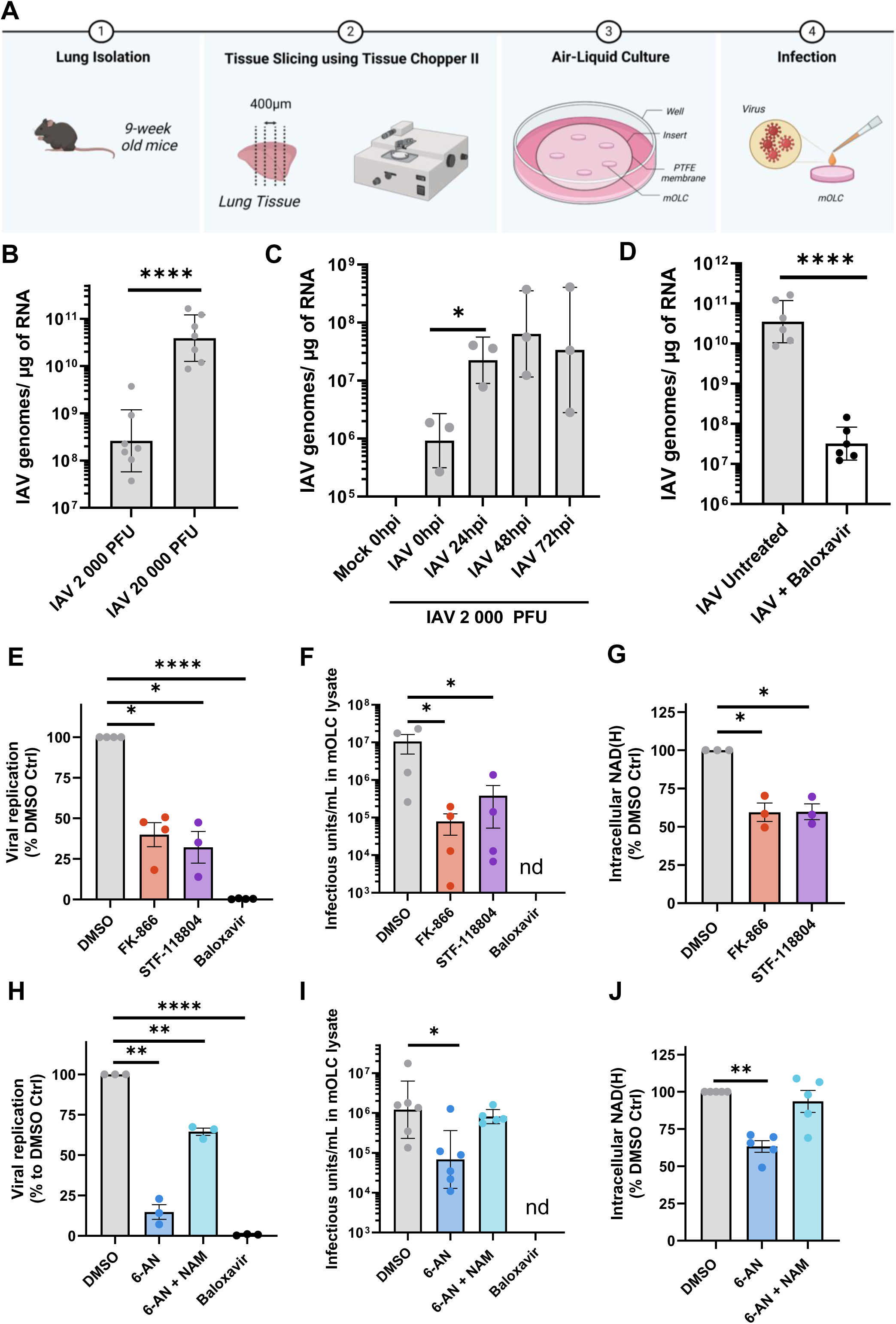
Inhibitors of NAD biosynthesis and metabolism restrain H3N2 replication and infectious viral particles production in murine organotypic lung cultures. (A) Procedure to generate murine organotypic lung cultures. After isolation, lung tissues were sliced at 400 µm thickness and cultured at the air-liquid interface using PTFE membrane inserts. mOLCs were infected by dropping viral inoculum on its surface. Treatments or DMSO solvent alone were added in the culture medium at the indicated concentration and mOLCs were treated by dropping drug-containing medium on its surface. (B-D) mOLCs were infected or not (mock) with 2 000 or 20 000 PFU of IAV before being treated or not with Baloxavir (100 nM). mOLCs were harvested just after (0), 24, 48 or 72 h post infection and viral M segment was quantified by RT-qPCR and normalized to murine Rplp0 expression. Data are presented as the geometric means ± SD of normalized IAV genome copy numbers / µg of RNA. (B) mOLCs were harvested at 48 hpi. Data from 7 independent animals. (C) Kinetic data from 3 independent animals. (D) mOLCs were harvested at 48 h post-infection with 20 000 PFU of IAV and treated or not with 100 nM Baloxavir. Data from 6 independent animals. (E-J) mOLCs were infected or not with 2 000 PFU of IAV before being treated with NAMPTi (20 µM for FK-866 or STF-118804), 6-AN (100 µM) with or without nicotinamide (500 µM), Baloxavir (100 nM) or DMSO solvent. Treatment of mOLCs as well as medium were renewed at 24 hpi. (E, H) Viral M RNA segment was quantified by RT-qPCR and normalized to murine Rplp0 mRNA. Data presented are means ± SEM of 3 experiments, each point is the pool of 3 mOLCs from different animals. (F, I) Concentration of viral infectious particles in mOLC lysates was determined by TCID_50_ assay on MDCK cells. Geometrical means ± SD of 4-6 independent animals is shown. (G, J) NAD(H) quantity measured in mOLC lysates. Means ± SEM of 3-5 different animals.

We then analyzed the impact of NAD pathway modulators, NAMPTi and 6-AN, on IAV replication in mOLCs. An inoculum of 2 000 PFU was used for the infection, and treatment was applied immediately after infection. Media and treatments were renewed at 24 hpi, and mOLCs and culture media were collected at 48 hpi. NAMPTi at 20 µM reduced IAV replication in mOLCs by approximately 60% as assessed by quantification of the M segment in the tissue (Fig. 3E). The production of infectious viral particles was measured by titration, showing a 2 log_10_ inhibition by FK-866 and a 1.5 log_10_ inhibition by STF-118804 (Fig. 3F). Baloxavir abolished IAV replication and viral production as expected (Fig. 3E-F). As expected, NAMPTi significantly reduced the pool of NAD by 40% in mOLC lysates (Fig. 3G). The antimetabolite 6-AN also reduced IAV replication by 90% (Fig. 3H), which is a more potent effect than observed in cell lines, and inhibited the production of viral infectious particles (Fig. 3I). The addition of NAM to 6-AN treatment partially restored IAV replication and production of infectious particles in the mOLC infection model (Fig. 3H-I). In these conditions, 6-AN reduced the pool of NAD in mOLCs by approximately 35%, and the addition of NAM restored NAD levels to 90% of the DMSO control (Fig. 3J). Providing increasing doses of NAM to lung culture resulted in elevated NAD levels compared to control without any impact on IAV replication (*SI appendix,* Fig. S3E-F), indicating that basal NAD content in mOLC is not a limiting factor of viral replication. The viability of mOLCs was evaluated by measuring intracellular RNA levels at 48 hpi and LDH release in the culture medium at 0, 24 and 48 hpi (*SI appendix*, Fig. S4A-D). Overall, FK-866 slightly reduced RNA levels and induced a minimal increase of LDH release after 48h in mOLC medium (*SI appendix*, Fig. S4A-B). Some cytokine markers of innate immunity were also monitored in the culture medium of mOLCs. Results showed that the secretion of IP-10 triggered by IAV was not affected by NAMPTi, while the production of IL-6 and MCP-1, which are mainly secreted by activated macrophages, was enhanced (*SI appendix*, Fig. S5A-C). This suggests that NAMPTi treatment favors lung macrophage activation. It is not the case of 6-AN, which inhibits the secretion of these cytokines and especially IP-10, whereas NAM tends to restore IP-10 induction by IAV (*SI appendix*, Fig. S5D-F). In summary, *ex vivo* results showed a significant decrease in IAV replication and viral production following NAMPTi or 6-AN treatment, and the addition of NAM reverted this effect, demonstrating IAV dependence on NAD.

### IAV replication in murine organotypic lung cultures results in NAD^+^ depletion and induction of NAD-consuming PARP enzymes

To better understand IAV dependence on NAD, we analyzed the metabolic modulations induced by IAV infection of mOLCs. Unsupervised semi-polar metabolomic analysis was performed at 48 hpi on mOLCs that were uninfected (mock), or infected with 20 000 PFU of IAV H3N2 strain and treated or not with Baloxavir (100 nM). Comparison between mock and IAV-infected slices identified 9 metabolites significantly modulated at 48 hpi (p-value<0.05 and fold change (FC) ≥1.2), one of which was identified as NAD^+^ (Fig. 4A). NAD^+^ levels were significantly reduced by 30% in the infected samples compared to the mock. Interestingly, NAD^+^ levels were restored to 90% of the mock level when IAV-infected slices were treated with Baloxavir (Fig. 4A). Another energy metabolism-related metabolite, adenosine monophosphate (AMP), was also modulated upon infection, showing a 25% increase that was prevented by Baloxavir treatment (*SI appendix*, Fig. S6). Pantothenic acid levels were reduced following IAV infection and partially restored upon Baloxavir treatment (*SI appendix*, Fig. S6). Since it is a precursor of coenzyme A (CoA) and acyl carrier protein (ACP), a decreased level of pantothenic acid in lung tissue indicates a potential defect in energy metabolism, surfactant synthesis, and antioxidant defense. Leucine and isoleucine levels were also significantly reduced, while pyroglutamic acid increased upon infection. However, their levels were not restored by Baloxavir treatment, suggesting that this modulation is independent of IAV replication (*SI appendix*, Fig. S6). Overall, metabolomics data showed that IAV infection results in metabolic alterations, and in particular NAD^+^ depletion in mOLCs.

**Figure 4.**
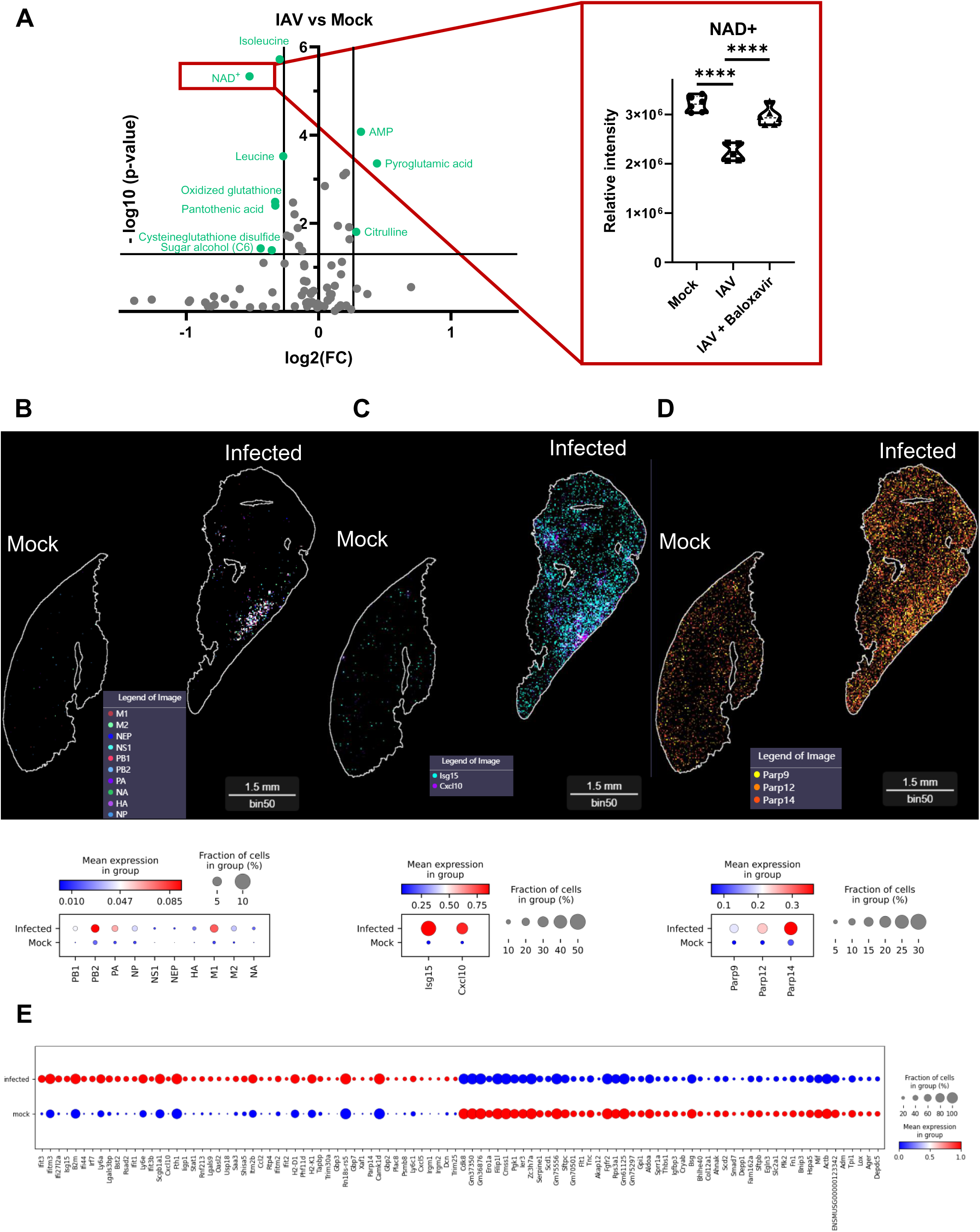
Enhanced NAD consumption in murine organotypic lung cultures upon IAV infection. (A) Relative abundance of metabolites at 48 hpi, expressed as relative peak intensity in mOLCs not infected (mock), infected with 20 000 PFU of IAV or infected and treated with 100 nM Baloxavir (IAV + Baloxavir). Two distinct pools of 5 mOLCs from independent mice were measured in triplicate for metabolite analysis by LC-MS/MS semi-quantitative analysis. (B-E) mOLCs from the same mice were infected or not with 20 000 PFU of IAV and incubated for 48 h. After fixation and inclusion in OCT, cryosections were prepared for spatial transcriptomic analysis. (B, C, D) Spatial view of mRNAs and dot plots presenting the intensity of expression and the fraction of positive cells for IAV proteins NP, M1/M2, HA, NA, NS1/NEP, PA, PB1, PB2 (B), mouse Cxcl10 and Isg15 (C) or Parp 14,12 and 9 (D). (E) Dot plots presenting the intensity of expression and the fraction of positive cells for the TOP 20 most differentially expressed genes between the infected versus the mock slice.

To further understand NAD^+^ depletion upon IAV infection, we performed high resolution Stereo-Seq spatial transcriptomic analysis on 12-µm thick cryosections of mOLCs that were cultured and infected as described above. We analyzed transcriptomic differences between infected and non-infected mOLCs at 25 µm resolution. The transcriptome sequences were aligned on the viral genome, and all viral mRNAs encoding IAV polymerase subunits (PA, PB1, PB2), structural proteins (NP, HA, NA, M1 and M2) and non-structural proteins (NS1 and NEP) were detected only in the infected mOLC (Fig. 4B). As expected, viral RNAs were mostly detected in a cluster enriched in pneumocytes detected by clustering of lung populations, based on the analysis of their transcriptomic signature (*SI appendix,* Fig. S7). We then analyzed the differentially expressed genes (DEGs) in the infected mOLC compared to the mock, determined with an adjusted p-value below 0.05. We detected a characteristic tissue response to the viral infection, with upregulation of ISGs such as Isg15 and Cxcl10 (IP-10) and enhanced expression of MHC class I molecules (H2-D1, H2-K1) (Fig. 4C-E). Three independent experiments were analyzed by

Stereo-Seq, highlighting a strong overlap in the modulated genes, with 99 DEGs conserved across experiments (*SI appendix,* Fig. S8A-D), including 53 genes that were strongly induced (log_2_(FC)>1.25) in at least one dataset (*SI appendix,* Fig. S9). The list of DEGs found at the intersection of the 3 experiments (*SI appendix,* Dataset S1) was analyzed for enrichment in biological processes (GO terms-BP). Genes involved in antiviral defense and interferon (IFN) response were overrepresented (*SI appendix,* Fig. S8E), reflecting the important induction of innate immunity. This is illustrated by Fig. 4C, which shows the spatial distribution of Isg15 and Cxcl10 expressions. Both genes were induced not only in the infected foci, but also around them, demonstrating that innate immunity is also activated in the vicinity of infected cells. Since NAD^+^ depletion in the infected mOLC could be explained either by the downregulation of NAD^+^ biosynthesis or by the upregulation of NAD^+^-consuming enzymes, we specifically analyzed the expression of genes involved in these pathways in infected vs mock-treated mOLC. As expected, NAMPT was highly expressed, whereas enzymes involved in *de novo* and Preiss-handler pathways of NAD biosynthesis (Fig. 1C) were expressed only at low levels (*SI appendix*, Fig. S10A). Their expression levels remained unchanged in the infected slices compared to the mock slices, except for NAMPT which was increased (*SI appendix*, Fig. S10A-C). We then analyzed the expression of genes encoding NAD^+^-consuming enzymes (*SI appendix*, Fig. S10D-F). Transcripts of Sarm1, CD38, Sirtuins 2, 7 and 11 were detected, but their expression levels remained unchanged across the 3 experiments (*SI appendix*, Fig. S10-E-F). Interestingly, several poly(ADP-ribosyl) polymerases (PARPs) were among the most significantly induced genes in the infected mOLCs, including Parp9, Parp12 and Parp14 (Fig. 4D, *SI appendix,* Fig. S9 and Dataset S1). We confirmed by RT-qPCR analysis their upregulation upon infection, whereas the expression of other NAD^+^-consuming enzymes such as Parp1, Parp2, Tankyrase 1 and 2, remained unchanged (Tnks, alias Parp5a and Tnks2, alias Parp5b) (*SI appendix,* Fig. S11). Parp9, 12 and 14 were not only induced in the infected foci, but also broadly around them like ISGs (Fig. 4D).

PARPs are ADP-ribosyl-transferases that use NAD^+^ as a substrate to add a mono- or poly-ADP-ribose unit to proteins, resulting in a post-translational modification (MARylation or PARylation, respectively). When added to proteins, these modifications alter their functions and interactions, and are therefore involved in various metabolic pathways and cellular responses to stress (25). PARP12 and PARP14 are MARylating PARPs, whereas PARP9 is catalytically inactive, but binds to PARP14 and modulates its activity. We used the inhibitor of MARylating PARPs, OUL232, to investigate the role of PARP activity, and found that it prevented NAD depletion upon IAV infection in mOLCs, suggesting that NAD consumption is linked to PARP induction (Fig.5A). OUL232 reduced viral production *in vitro* in A549 infected cells (*SI appendix*, Fig. S12A). Western blot analysis revealed that both OUL232 and FK-866 reduced global protein MARylation in IAV-infected cells, consistent with the involvement of NAD-dependent ADP-ribosylation pathways (*SI appendix*, Fig. S12B-D). *Ex vivo*, OUL232 also reduced IAV genome replication in a dose-dependent manner at 2 and 10 µM (Fig. 5B), and the production of infectious viral particles in mOLC lysates was decreased by approximately 1 log_10_ (90%) (Fig. 5C), while other inhibitors OUL35 and PARP10/15-IN1, mainly targeting PARP10 in mice or H-10, an inhibitor of PARP14, had no significant impact in these conditions (*SI appendix*, Fig. S13A-C). In line with previous studies showing that PARP1 and TNKS1/2 proteins are needed for IAV replication (26–28) and an in vivo antiviral assay (29), PARP1/2 inhibitor Olaparib and TNKS inhibitor Nesuparib reduced infectious viral particles production (*SI appendix*, Fig. S13A,D). All PARP inhibitors showed no cytotoxicity on mOLCs at the concentrations used as assessed by RNA quantification and LDH release (*SI appendix*, Fig. S14A-F). The combination of OUL232 and FK-866 exerted a potent antiviral effect, reducing IAV production by 2 log_10_ (Fig. 5D). At this high concentration of FK-866, the addition of OUL232 did not significantly further reduce infectious viral production, suggesting that their antiviral effects may involve overlapping NAD-dependent processes. Collectively, these results show that pharmacological targeting of NAD biosynthesis and PARP activities reduces IAV replication and infectious viral production, highlighting the contribution of NAD-dependent pathways to IAV infection (Fig. 5E).

**Figure 5.**
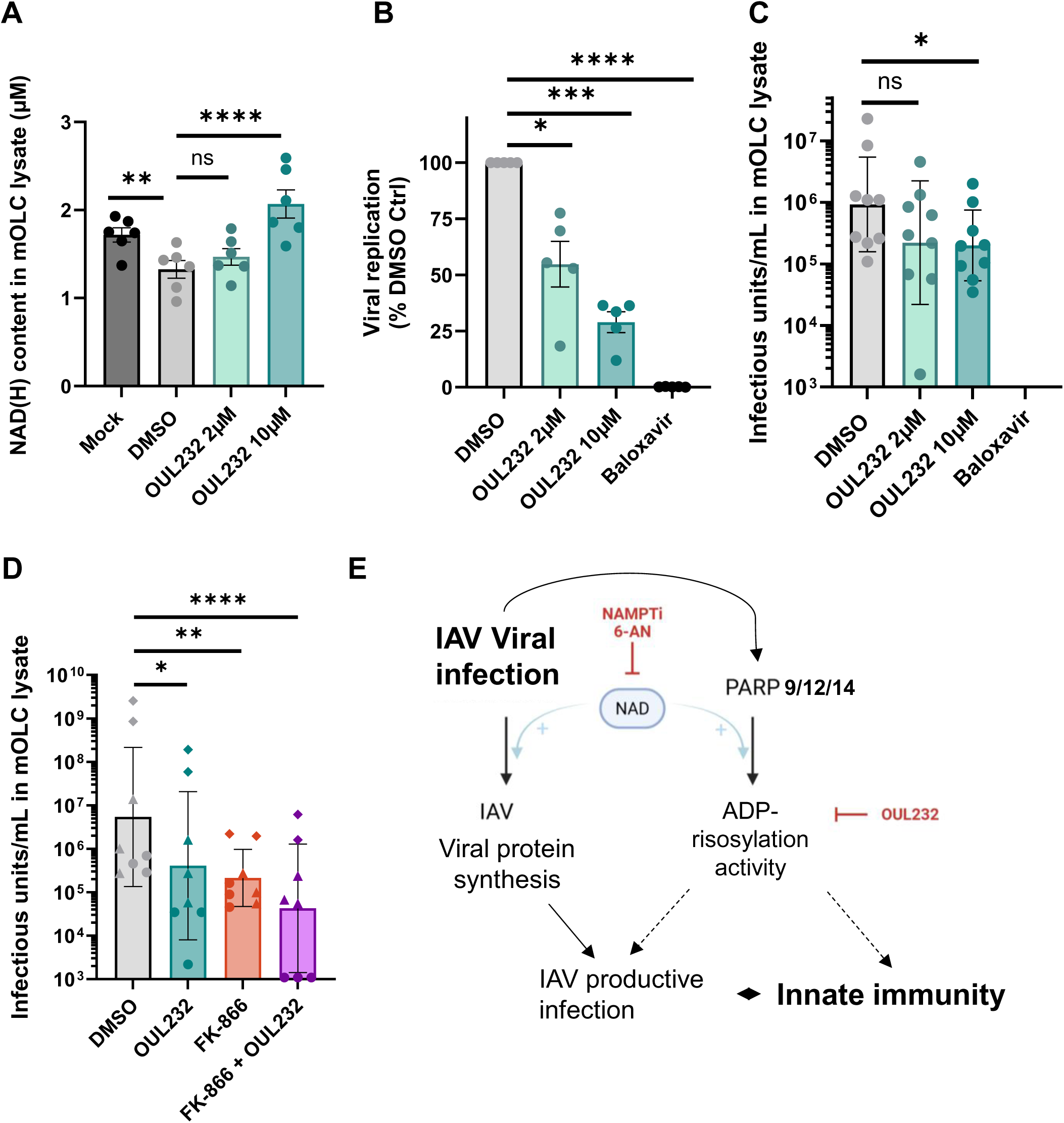
NAD-dependent ADP-ribosylation promotes viral infection in mOLCs. mOLCs were infected with 20 000 PFU (A) or 2 000 PFU (B,C) IAV before being treated with the indicated drug or DMSO solvent. Treatment of mOLCs as well as medium were renewed 24 h post-infection. Medium and tissues were collected at 48 hpi. (A) Three slices from 3 animals were pooled and lysed in buffer before NAD(H) assay. (B) Three slices from 3 different animals were collected and pooled for RNA extraction. Viral M RNA segment was quantified by RT-qPCR and normalized to murine Rplp0 mRNA. Data presented are the mean ± SEM of 4 experiments, each point is the pool of 3 mOLCs from different animals. (C) Concentration of viral infectious particles in mOLC lysates was determined by TCID_50_ assay on MDCK cells. Geometrical means ± SD of 9 independent animals is shown. (D) mOLCs were infected with 20 000 PFU before being treated with 10 µM OUL232 or 20 µM FK-866 or both. Concentration of viral infectious particles in mOLC lysates was determined by TCID50 assay on MDCK cells. Geometrical means ± SD of 8 independent animals is shown. Distinct infection experiments are indicated by distinct symbol shapes. P-values were obtained from paired ANOVA tests. (E) Role of NAD in IAV viral life cycle. As a metabolic cofactor, NAD fuels IAV biosynthetic processes and NAD is metabolized by PARPs to ADP-ribosylate target proteins, modulating innate immunity and IAV replication.

## Discussion

This study highlights the dependence of IAV replication on NAD, which serves both as a cofactor for numerous redox reactions and as a substrate for NAD-consuming enzymes (30, 31). Through *in vitro* screening of a library of metabolic regulators, we found that two chemically distinct NAMPTi, FK-866 and STF-118804, exhibit antiviral activity against IAV (Fig. 1). The antiviral activity of FK-866, observed here in the nanomolar range, extends recent findings obtained at micromolar doses (32). We next established an *ex vivo* model of IAV infection in adult mouse lung tissue, preserving tissue architecture and resident immune cells while enabling the characterization of early local immune responses to viral infection. Using this model, we showed that depleting the NAD pool with NAMPTi or interfering with NAD metabolism using 6-AN (Fig. 2) both limit IAV replication and infectious particle production, whereas supplementation with an excess of NAD precursor restores these parameters (Fig. 3). Furthermore, metabolomic analyses revealed that IAV infection leads to the depletion of NAD^+^ in the infected tissue, while spatial transcriptomic analysis showed the induction of MARylating PARPs, especially PARP12 and PARP14 that consume NAD^+^ to ADP-ribosylate their targets (Fig. 4). Accordingly, inhibition of MARylating PARPs with OUL232 prevented NAD depletion triggered by IAV infection and reduced IAV productive infection in the tissue (Fig. 5A-C). Combined treatments by FK-866 and OUL232 suggest that both molecules are acting on the same pathway to restrain IAV infection of the lung (Fig. 5D). Taken together, these results suggest that NAD serves both as an essential cofactor that fuels cellular metabolism and as a substrate for MARylating PARPs upon which the virus depends for its replication and/or evasion of the innate immunity (Fig. 5E). This identifies a potential vulnerability point that could be exploited to develop antiviral strategies.

### Targeting NAD biosynthesis as a host-directed antiviral strategy

Lowering the NAD pool to restrain viral replication has long been suggested as a host-directed antiviral strategy. The antimetabolite 6-AN was first described in 1964 as inhibiting Vaccinia virus production in chick-embryo fibroblasts (33) and more recent studies have established the antiviral effect of NAD metabolism inhibitors against Zika, Hepatitis B, Hepatitis C, and Dengue viruses (18–20). The antiviral activity of 6-AN correlated with a decrease in the NAD^+^ pool and was reversed by NAD precursor supplementation (18, 19). Chan et al. similarly found that 6-AN reduced NADPH production and diminished IAV replication in MDCK cells, that they attributed to inhibition of glucose-6-phosphate dehydrogenase (G6PD), restraining the activity of the pentose phosphate pathway (21). As a nicotinamide analog, 6-AN is converted by NAD kinase into 6-amino-NAD and 6-amino-NADP, which inhibit numerous NAD(P)-dependent enzymes, notably 6-phosphogluconate dehydrogenase (6PGD) and G6PD in the oxidative pentose phosphate pathway. The precise enzymatic requirements for IAV replication, however, remain to be defined. Consistent with earlier studies on liver cells (22), we found that 6-AN strongly reduced the glycolytic activity of lung epithelial cells while NAM supplementation restored it to control level (Fig. 2). The NAMPT inhibitor FK-866 led to a pronounced depletion of the NAD pool by acting as a direct inhibitor of the salvage pathway (Fig. 1D). This resulted in reduced glycolytic activity and respiration of cells *in vitro* (Fig. 2). In the *ex vivo* model, we showed that FK-866, STF-118804 and 6-AN all led to a reduction in the available NAD pool, which was associated with antiviral effects. In line with these observations, Lv *et al.* recently reported that oral administration of 10 mg/kg/day of FK-866 before and during infection, restrained IAV H1N1 PR8 infection of mice (32). However, the pre-treatment used in this study highlights the need to improve the experimental strategy.

The NAD pathway first emerged as a target in oncology, as tumor cells consume large amounts of NAD to sustain their high proliferation rate (34). Targeting NAD metabolic pathways for a long period of time, in an attempt to control tumor progression, has raised concerns about the toxicity or side effects of 6-AN or NAMPT inhibitors (34–36). However, short term treatments during acute viral infections could be more sustainable. Moreover, promising strategies are emerging to mitigate these concerns, including targeted NAMPT degradation and antibody-drug conjugates that deliver NAMPT inhibitors directly to the targeted tissue. In the lung, most of the NAD is synthesized from NAM via the salvage pathway, involving NAMPT and nicotinamide nucleotide adenylyltransferase (NMNAT) enzymes (Fig. 1C) (30, 37). In agreement, our spatial transcriptomic data revealed high abundance of NAMPT transcripts with expression of NMNAT isoforms 1-3 in both infected and uninfected mOLCs. In contrast, the initiating enzymes of the *de novo* and Preiss-Handler pathways (Fig. 1C), the nicotinate phosphoribosyltransferase (NAPRT) and the quinolinate phosphoribosyltransferase (QPRT) respectively, are weakly expressed (*SI appendix*, Fig. S10A-C). Thus, targeting NAMPT directly into the lungs is of prime interest to modulate NAD-dependent processes in this organ. NAMPT inhibition may also deplete the NADP pool, even though Liu et al. highlighted a particularly low NAD kinase forward flux in murine lungs, with only 10% of produced NAD being phosphorylated into NADP (38). Thus, in the context of severe respiratory infection, transient and lung-targeted NAMPT inhibition could represent a promising strategy to limit viral replication by reducing NAD availability. However, its therapeutic potential will require further in vivo evaluation, particularly regarding treatment tolerability and its impact on pulmonary inflammation. As a host-directed approach, this strategy also holds pan-viral potential against emerging viruses.

Interestingly, our results also revealed a differential impact of NAMPTi and 6-AN on the innate immune response to IAV infection. Although NAMPTi prevented the induction of cytokines such as IP-10 triggered by IAV infection *in vitro* (Fig. 1G–H), the secretion of this antiviral cytokine was maintained in the infected lung tissue upon treatment by NAMPTi and PARP inhibitor OUL232 (*SI appendix,* Fig. S5). Moreover, IL-6 and MCP-1 secretions which are indicative of macrophage activation, were enhanced or conserved, supporting the activation of innate immunity in these conditions. In contrast, 6-AN broadly suppressed all these cytokines secretion *ex vivo* (*SI appendix,* Fig. S5). *In vivo*, Lv et al. observed that FK-866 pre-treatment attenuated H1N1-induced inflammation in mice, with reduced TNF-α, IL-6, and IFN-α levels in the lungs at days 3 and 5 post-infection (32). Future studies will need to evaluate whether reshaping the inflammatory response, along with inhibiting the replication of IAV, is beneficial when using inhibitors of the NAD pathway as a host-targeting antiviral strategy.

### IAV infection, PARP induction, and NAD-consuming MARylation

Modulation of the NAD pool has already been documented during diverse viral infections. Grady et al. showed a rapid and drastic NAD^+^ depletion upon Herpes Simplex Virus 1 of human fibroblasts (39). *In vivo*, patients with severe COVID-19 displayed significantly lower circulating NAD^+^ than healthy individuals. Additionally, increased transcription of salvage pathway enzymes as observed during SARS-CoV-2 infection both *in vitro* and *in vivo* in mice and humans (40–42). It has also been shown that depletion of the NAD pool underlies Zika virus-induced microcephaly (43). Here, we show that IAV infection also leads to the depletion of NAD in the lung tissue when using the *ex vivo* mOLC model. Interestingly, both 6-AN and FK-866 had a stronger impact on IAV in the mOLC model than in A549 cells, resulting in a more pronounced reduction in infectious particle production. This may reflect the higher sensitivity of the diverse primary cells within the lung tissue to NAD level modulation (44, 45), as well as the greater physiological relevance of low infectious doses in *ex vivo* experiments.

Our spatial transcriptomic results further revealed that Parp14, Parp12 and Parp9 transcription was increased in infected mOLCs, with maximal expression in infected areas (Fig. 4D), a pattern consistent across all three analyzed samples (*SI appendix*, Fig. S9). PARPs are mono- or poly-ADP-ribosyl-transferases, resulting in a post-translational modification (MARylation or PARylation) of cellular or viral target proteins. An increased PARylation activity triggered by viral infection has been documented for several viruses, alongside with the induction of PARP expression, and these modifications can either restrict or promote viral replication (42, 46). They are reversible, and some viral proteins are able to remove the ADP-ribose moiety, thereby limiting PARP activity (47). The PARP family comprises 17 human and 16 mouse members, with distinct structures, localizations, enzymatic activities, and RNA-binding capacities (48). Whereas viral infection and IFN signaling can trigger the expression of some PARPs that contribute to antiviral host response (42), other studies indicate that IAV takes advantage of PARP expression (27, 49–51). Notably, the administration of PARP1/2 inhibitor Olaparib reduced IAV-induced mortality, lung injury and cytokine production in mice (29). Interestingly, it has been shown that IAV replication complex associates with PARP1, whereas Lv et al. recently reported that NAMPT plays a structural role in PB1 interaction with PA (32). Whether the proximity of NAMPT and PARP1 within the viral replication complex favors the formation of a metabolon-like complex that produces and consumes NAD to facilitate IAV replication remains to be determined.

Several MARylating PARPs are directly linked to innate immunity. Hence, some PARPs, especially PARP12, are ISGs or increase the expression of ISGs, like PARP9, or contribute to IFN-β secretion induced by pathogens, like PARP14 (31, 52, 53). These genes were also induced upon SARS-CoV-2 infection in human lung cell lines and *in vivo* in ferret tracheas, mice lungs, and human bronchoalveolar lavages (41, 42). Mining public transcriptomic data (GSE206606), we observed that PARP9, 10, 12 and 14 were significantly induced upon IAV H3N2 infection of A549 cells and also of primary human alveolar epithelial type II cells (54). We therefore used OUL232, an inhibitor of MARylating PARPs (PARP7, 10, 11, 12, 14, and 15), which based on published IC50 parameters, should preferentially inhibit PARP10, 7, 12, 11 and 14, in this order (55) (*SI appendix* Fig. S13A). OUL232 treatment prevented NAD depletion triggered by IAV infection and reduced both viral replication and infectious particle production in infected mOLCs (Fig. 5A-C), revealing a global pro-viral role of mono-ART activity expressed in mOLCs upon IAV infection. The absence of a significant additive antiviral effect when OUL232 was combined with high doses of FK-866 suggests that both compounds converge on the same NAD-dependent pathway: by depleting the NAD pool, FK-866 likely abolishes the activity of the NAD-consuming enzyme targeted by OUL232, rendering its direct inhibition redundant (Fig. 5E). This supports the idea that the antiviral effect of OUL232 relies on a NAD-dependent enzymatic process, rather than on an independent or parallel mechanism.

The lack of highly selective inhibitors for individual PARPs remains a significant challenge in defining the exact enzymatic drivers of this process. In contrast to OUL232 pan-MARylating inhibitor, selective inhibitors of PARP10 (55–57) and PARP14 (58, 59) failed to reduce viral infection in mOLCs. Olaparib also showed a slight inhibitory effect on viral replication in our model (*SI appendix,* Fig. S13). It remains to be determined whether this effect stems from Olaparib’s ability to block PARP1/2 or other MARylating PARPs (56, 60, 61). Given that PARP12 enzyme activity can negatively regulate the IFN response, and its invalidation partially protected mice from IAV challenge (62), PARP12 represents a plausible candidate for the pro-viral activity observed here. However, deciphering whether these effects are driven by PARP12 or a combination of multiple PARPs will require targeted investigations in future studies.

Finally, preventing virus-induced metabolic remodeling can restrain viral replication and our results indicate that targeting NAD metabolic pathway could be a valuable strategy for developing host-targeting antiviral directed against IAV infection. Indeed, targeting the host rather than the virus itself as a therapeutic approach offers a number of advantages, notably overcoming the genetic diversity between IAV strains and limiting the risk of the emergence of resistant variants.

## Materials and Methods

### Viruses

Mouse adapted H3N2 Influenza A/Scotland/20/74 strain was amplified on Madin-Darby Canin Kidney (MDCK) cells as previously described (63). Viral stocks used within this study were titrated by Plaque Assay on the same cells, aliquoted and stored at -80°C. Freshly thawed viral stocks were diluted in the cell culture medium used in the assays described below.

### Cells and Media

A549 cells were cultured in DMEM, high glucose with GlutaMAX (10566016; Gibco). BEAS-2B cells were grown in DMEM/F-12K Medium, high glucose with GlutaMAX (10565018; Gibco), and MDCK cells were grown in MEM medium, high glucose with GlutaMAX (41090036; Gibco), all supplemented with 10% fetal calf serum (FCS) (FB-1285; Biosera) and 100 U/mL penicillin/streptomycin (15140-122; Gibco). All cells were incubated at 37°C under 5% CO2.

### Reagents

Following reagents were purchased from MedChemExpress: 6-aminonicotinamide (6-AN - HY-W010342), nicotinamide (NAM - HY-B0150), OUL232 (HY-148566), FK-866 (HY-50876), STF-118804 (HY-12808), Baloxavir (HY-109025A). They were dissolved in DMSO except for 6-AN and NAM, which were dissolved in sterile apyrogenic water (600500; Aguettant). The same amount of solvent was added in the untreated control (DMSO).

### Treatment and infection of *in vitro* models

For experiments with pretreatment, cells were seeded in standard culture medium and treated with the antiviral molecule to reach the indicated final concentration. 24 h later, cells were infected by the addition of H3N2 Influenza A/Scotland/20/74 at the indicated Multiplicity Of Infection (MOI). Cells were incubated for a further 24 or 48 h, and the culture supernatants and cell lysates were harvested and stored at -80°C until analysis.

For experiments with treatment post-infection, cells were seeded 24 h before infection. They were washed twice with PBS and infected for 1.5 h at 37°C with H3N2 Influenza A/Scotland/20/74 at the indicated MOI in standard culture medium devoid of FCS with 1 µg/mL TPCK-treated Trypsin (T1426-MG; Sigma). Then, cells were washed once with PBS and incubated with treatment at the indicated final concentration in culture medium with 0.2 µg/mL TPCK-treated trypsin, without FCS for A549 or with 10% FCS for BEAS-2B cells. Supernatants and cell lysates were collected 24 hpi and stored at -80°C before analysis.

### Screening of metabolic regulators library

The DiscoveryProbe Metabolism-related Compound Library from APExBio (L1032), containing 493 compounds dissolved in DMSO (unless otherwise specified), was screened at a final concentration of 10 µM in 96-well plates. Briefly, A549 cells (30 000/well) were seeded in wells containing metabolism related compounds one day before infection with H3N2 Influenza A/Scotland/20/74 strain at MOI 0.5. On each plate, DMSO-treated infected cells were used as positive control (4 wells/plate) and Baloxavir (20 nM) was used as a reference antiviral treatment (4 wells/plate). Non-infected cells were placed as negative controls in the last plate of the screening (8 wells). At 24 and 48 hpi, 25 µL of supernatants were collected for neuraminidase activity (NA) assay. NA fluorescent signal measured in each well was normalized by the average fluorescence intensity of its plate. Nuclei were labeled for 45 min at 37°C by addition of Hoechst 33342 (20 µM final concentration) at 48 hpi and fluorescent signal was monitored with a TECAN infinite M200 instrument to evaluate cell viability. The Z’ coefficient was calculated using the NA signal from DMSO-treated and Baloxavir-treated wells to evaluate the overall quality of the screening procedure (16). A Z′ factor below 0.5 (0 < Z′ < 0.5) indicates some variability in the results or a limited dynamic range, compared with an ideal assay where Z′ > 0.5.

### Neuraminidase activity assay

Standard fluorometric assay was used to measure Influenza virus neuraminidase activity. In a black 96-well plate, 25 µl of cell supernatants were diluted in 25 µl of D-PBS containing calcium and magnesium. The reaction was started with 50 µl of 20 mM 2’-(4-Methylumbelliferyl)-a-D-N-acetylneuraminic acid sodium salt hydrate (4-MUNANA, M8639; Sigma). After 1 h incubation at 37°C, the reaction was stopped by adding 100 µl of 0.1 M glycine pH 10.7 / 25% ethanol. The fluorescent cleavage product was quantified with a Tristar 5 microplate reader (Berthold) at 365 nm excitation and 450 nm emission wavelengths.

### Cell count measurement

To assess the effect of the drugs on cell count, cells were treated as described in the figure legend. After 24 h of treatment, nuclei were labeled with a 20 µM Hoechst 33342 (62249, Thermo Fisher Scientific) solution for 45 min at 37°C before automated cell counting with the microplate-based Imaging cytometer Celigo (Revvity).

### Quantification of viral protein expression

Viral nucleoprotein (NP) and non-structural protein (NS1) labeling was carried out on cells fixed with 4% formaldehyde for 20 min and permeabilized with 0.5% Triton X-100. Anti-NP staining was performed using the rabbit polyclonal anti-Influenza NP antibody (PA5-32242; Thermo Fisher Scientific) at 1:400, anti-NS1 staining was performed using the rabbit polyclonal anti-Influenza NS1 antibody (GTX125990; Genetex) at 1:500, both in combination with goat anti-rabbit Alexa Fluor 488 (AF488) antibody (A-11034; Thermo Fisher Scientific) at 1:1000. Total cell count was assessed by nuclei labeling with Hoechst 33342 reagent as above. Images were acquired using the Celigo microplate-based multichannel imaging cytometer (Revvity). For each condition, mean AF488 fluorescence intensity was quantified from three images per well, across triplicate wells, using ImageJ software. Background signal was assessed from mock-infected wells and subtracted accordingly. For each independent experiment, AF488 signal was normalized to the DMSO-treated infected control.

### Cells glycolysis and respiration monitoring

Cells were seeded in poly-L-Lysine-coated, XF 24-wells cell culture microplates (Agilent) in 125 μL cell culture medium. After 4 h, cells were treated for 24 or 48 h with 80 nM FK-866, 100 µM 6-AN, with or without 500 µM NAM or the solvent alone by addition of 125 µL of drug-containing medium. The Glycolytic Rate assay (Agilent) was performed in 500 μL of Seahorse assay medium (XF DMEM pH 7.4 + 10 mM Glucose, 2 mM Glutamine, and 1 mM sodium pyruvate) following the manufacturer’s recommendations. The Proton Efflux Rate (PER) and Oxygen Consumption Rate (OCR) were measured using the Extracellular Flux Analyzer (XFe24 Seahorse, Agilent) at the basal stage, after inhibition of respiration by 1 μM Rotenone + Antimycin A, after injection of 50 mM of glycolytic inhibitor 2-deoxyglucose (2-DG). Cell numbers were determined post-assay by Hoechst staining and counted using the Cytation 1 image reader (Agilent). PER and OCR values were normalized to cell number.

### Animals and handling

Eight-week-old C57Bl/6 female mice were obtained from Janvier Labs (Le Genest Saint-Isle, France) with a clean health monitoring report. Mice were housed for 1 week with ad libitum food and water before being sacrificed for *ex vivo* experiments.

### Generation of murine organotypic lung cultures

Murine organotypic lung cultures (mOLCs) were generated as previously described with minor modifications (23, 24). Briefly, lungs from sacrificed 9-week-old adult mice were isolated and sliced using McIlwain Tissue Chopper II (Campden Instrument) at 400 µm thickness. Slices were kept in HibernateTM-A medium (A12475-01; Gibco) with 1% P/S, before being transferred on PTFE cell culture insert (PICMORG50; Millipore). mOLCs were cultured at the air-liquid interface in MEM GlutaMAX medium with HEPES and Earle’s salts (42360-024; Gibco) supplemented with 5 g/L D-glucose (G7528; Sigma), 0.1 mg/L human recombinant insulin (I9278; Sigma) and 25% horse inactivated serum (26050-088; Gibco).

### Infection and treatment of mOLC

mOLCs were infected with either 2 000 or 20 000 PFU of H3N2 IAV by depositing on the slice 2.8 µL of viral suspension diluted in MEM medium without serum. Mock mOLCs received the same volume of MEM medium. The treatment was introduced in the culture medium at the indicated final concentration, and 3 µL of culture medium were directly added on slices. Medium and treatment were renewed every 24 h unless otherwise specified.

### NAD(H) quantification

Total NAD(H), which corresponds to both NAD^+^ and NADH, was quantified using the NAD/NADH-Glo assay (G9071, Promega) or NAD^+^/NADH assay kit (MAK460, Sigma) according to manufacturer instructions. For *in vitro* experiments, the culture medium was removed and replaced by PBS before addition of NAD/NADH-Glo detection reagent (v:v). After 30 min incubation at 37°C, the luminescence signal was measured using a Tristar 5 microplate reader (Berthold). For *ex vivo* experiments, mOLC were dry frozen and stored at -80°C until NAD(H) quantification. The day of the assay, mOLCs were transferred into Precellys tubes containing silica beads (P000918-LYSK0-A; Bertin) with 200 µL of cold PBS. mOLCs were lysed twice using a Precellys Evolution homogenizer (Bertin) with the following protocol: 4x (10 s 6600 rpm – 10 s break). Lysates were then clarified at 17,000 g, 10 min at 4°C before NAD/NADH levels quantification as described above.

### RNA extraction and RT-qPCR

Total RNA was isolated from cells using the NucleoSpin RNA isolation kit (740955-250; Macherey Nagel) according to the manufacturer’s instructions. For *ex vivo* experiments, mOLCs were lysed in 500 µL of Tri Reagent (TR118-200; Molecular Research Center) in Precellys tubes with two steel beads (SSB23; Next Advance) with the Precellys Evolution homogenizer protocol: 3x (15 s 6,500 rpm – 15 s break). RNA was isolated from mOLC lysates using Direct-zol RNA Microprep (R2060; Ozyme) according to the manufacturer’s instructions.

RNA was reverse transcribed into cDNA using the High-Capacity RNA to cDNA (4387406; Applied Biosystems) kit with random primers and oligo dT. Quantitative PCR was carried out on cDNA templates diluted 10 times, with Quantinova® SYBR® Green PCR (208056, Qiagen) kit on a StepOnePlus Real-Time PCR System (Applied Biosystems), using 10 µM of the specific primers described in Table I. Results were analyzed with StepOne version 2.3 (Applied Biosystems). Gene expression was quantified using the 2^-ΔΔCt^ method. Calculations of the copy numbers were normalized to the standard deviation (SD) of the housekeeping gene (Rplp0) mRNA.

**Table I.**
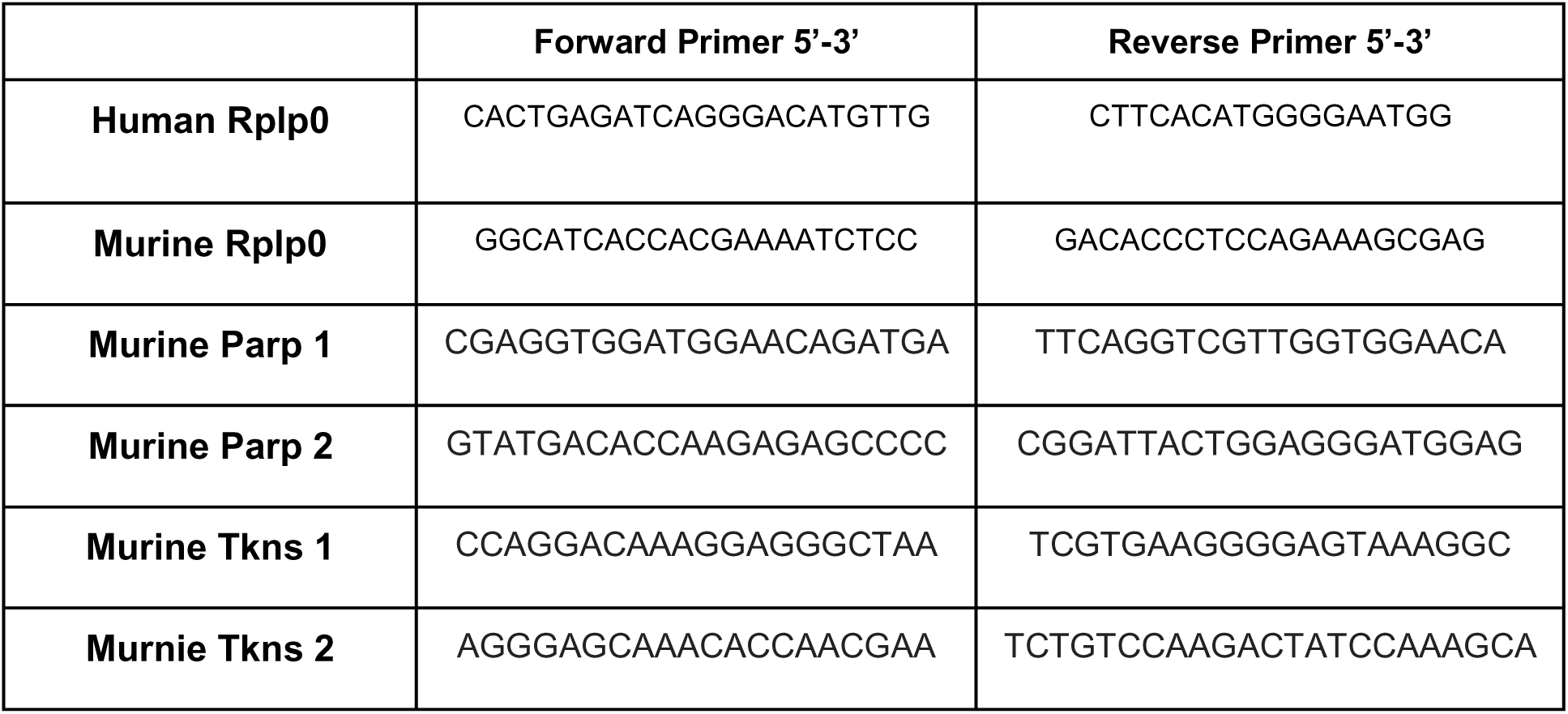
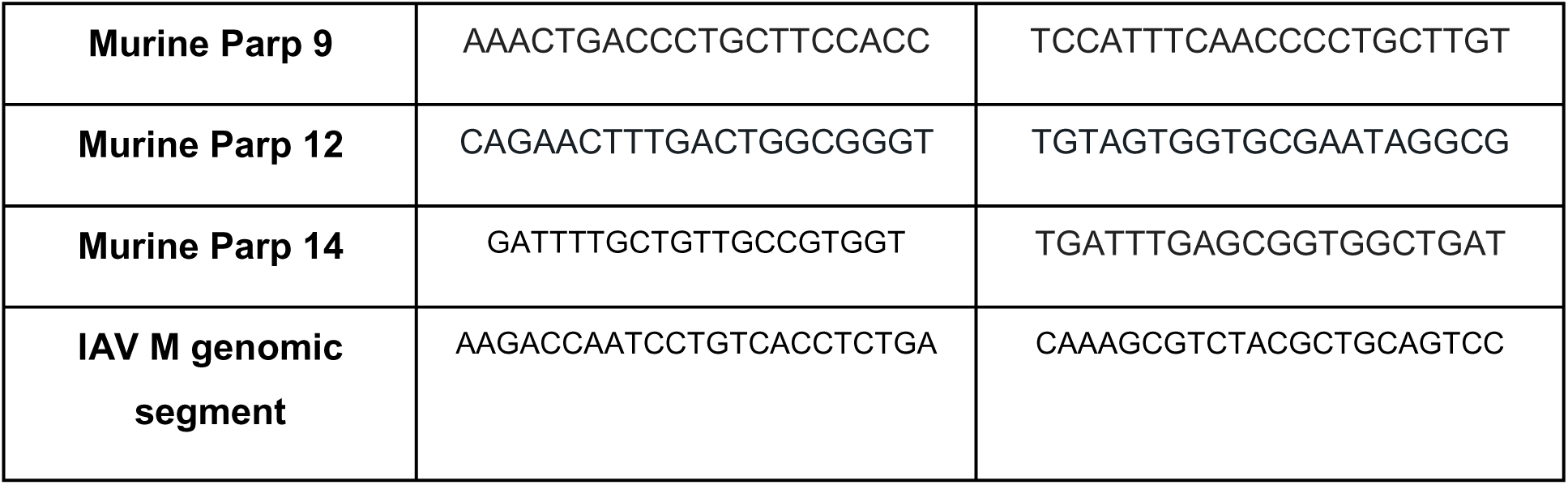
Sequence of the primers used in this study.

### IAV titration by TCID_50_ Assay

For *in vitro* experiments: harvested supernatants were thawed and serially diluted (10-fold) in MDCK MEM medium without FCS, supplemented with 1 µg/mL TPCK-treated trypsin, referred to as MDCK infection medium.

For *ex vivo* experiments: mOLCs were collected in 240 µL cold MEM GlutaMAX medium with 1% P/S and lysed twice using the Precellys protocol: 3x (15 s 6500 rpm – 15 s break) before being stored at -80°C. The day of the titration, mOLC lysates were thawed and clarified 10 min, 14,000 g at 4°C, before being serially diluted in MDCK infection medium as above.

MDCK cells were seeded 2 days prior to the assay, washed with PBS before inoculation with each dilution (8 replicates per dilution). Cells were incubated for 2 days at 37°C under 5% CO2. Cells were fixed and stained with 0.3% crystal violet - 20% methanol for 20 min. The infectious viral titer (infectious unit IU/mL) was determined by observation of cytopathic effect, using the Reed and Muench statistical method.

### Cytokines secretion assay

Culture supernatants were collected, clarified and stored at -80°C. The cytokines Interleukin (IL)-8, IL-6, and interferon gamma-induced protein 10 (IP-10) were quantified using cytokine-specific Cytometric Bead Array (CBA) Flex Sets (BD Biosciences) and analyzed on an LSR Fortessa II flow cytometer (BD Biosciences). CBA Flex Sets were obtained from BD Biosciences: IP-10 (558280), IL-6 (558276), IL-8 (558277), along with the CBA Master Buffer Kit (558265). U-PLEX Biomarker Group 1 (mouse) was also used for IL-6 monitoring using electro-chemiluminescent labels (MSD GOLD SULFO TAG) on U-PLEX assay platform by Meso Scale Discovery (MSD, Rockville, USA).

### Quantification of mono-ADP ribose by Simple Western

0.15 x 10^6^ A549 cells were seeded in 24-well plates with 0.5 mL of culture medium containing 10 µM OUL232, 80 nM FK-866, both OUL232 and FK-866 or their DMSO solvent alone. After incubation for 24 h, cells were infected with H3N2 Influenza A/Scotland/20/74 at MOI 0.1 for 2 h, then treated with 2 µM proteasome inhibitor MG-132 (M7449; Sigma-Aldrich). At 24 hpi, cells were washed with cold PBS and lysed on ice with 0.1 mL MOPS-RIPA 1X lysis buffer (20-188; Millipore; France) supplemented with Protease Cocktail Inhibitor (P8340; Sigma-Aldrich), 1 µM PDD17273 (SML1781; Sigma-Aldrich) and 40 µM PJ-34 (528150; Sigma-Aldrich). Lysates were collected, sonicated and clarified by centrifugation at 4°C (10 min; 17000 g). After protein quantification (DC Protein Assay; Bio-Rad; France), mono-ADP-ribosylated proteins were quantified with the Jess Simple Western System (Bio-Techne) following manufacturer’s instructions (12-230 kDa Fluorescence Separation Module, SM-FL001; EZ Standard Pack 1; PS-ST01EZ; Total Protein Detection Module for Chemiluminescence based total protein assays, DM-TP01; RePlex™ Module, RP-001). Protein extracts were dispensed in the assay plate at a final concentration of 1 mg/mL. Mono-ADP-ribosylated proteins were detected with a rabbit monoclonal antibody (HCA354; 1/10 final dilution; Biorad) and quantified by integration of the area under curve with the Compass for Simple Western Software.

### Semi-polar metabolomic analysis

All Precellys and Eppendorf tubes were pre-rinsed with 80% MS-grade MeOH (900688; Sigma-Aldrich), prepared with sterile apyrogenic water (600500; Aguettant, France). Five independent mOLCs were pooled and lysed in 1500 µL ice-cold 80% MS-Grade MeOH supplemented with 60 µM ^13^C6 L-Arginine (2061.1; Roth Sochiel) internal standard, using Precellys apparatus in tubes containing 1.4- and 2.8-mm silica beads (P000918-LYSK0-A; Bertin), with the following protocol repeated 3 times: 4x (10 s 6600 rpm – 10 s break). mOLC lysates were collected into Eppendorf tubes. Precellys tubes were washed with 1500 µL of ice-cold 80% MeOH with ^13^C6 L-Arginine, that was also collected. These lysates and washes were clarified at 17,000 g for 5 min at 4°C, pooled for each condition and analyzed in triplicate after storage at -80°C.

Samples preparation and analysis were performed by cmbio (Denmark) with a method described by Doneanu et al. slightly modified (64). Before analysis, each sample was diluted 11 times in mobile phase A eluent (10 mM ammonium formate + 0.1% formic acid in ultrapure water) and fortified with stable isotope labeled standards. Analysis was carried out using a UPLC system (Vanquish, Thermo Fisher Scientific) and a high-resolution quadrupole-orbitrap mass spectrophotometer (Orbitrap Exploris 240 MS, Thermo Fisher Scientific). The ionization was achieved with an electrospray ionization interface operated in positive and negative ionization mode under polarity switching. A QC sample was analyzed in MS/MS fragmentation mode for the identification of compounds. Data were processed by Compound Discoverer 3.3 (Thermo Fisher Scientific) and Skyline 23.1 (65).

A total of 1,347 compounds were detected in the samples. Hereof, 130 metabolites could be identified with enough confidence; 33 were annotated by accurate mass, MS/MS spectra and known retention time (level 1) and 97 were annotated by accurate mass and retention time (level 2a) obtained from standards analyzed on the same system. The relative abundance of each annotated metabolite (levels 1 and 2a) was calculated. Univariate statistics (provided as t-tests and relative intensity fold changes) show if any single variable is significantly different between two specified treatment groups. Metabolites were considered significantly regulated when fold change was varying by more than 20% and the p-value was less than 0.05.

### Immunofluorescent staining of mOLCs

mOLCs were collected, fixed for 2 h in 4% paraformaldehyde (PFA; 047340.9M, Thermo Fisher Scientific) and washed 3 times in PBS before quenching in PBS - 0.1% glycine. mOLCs were then permeabilized overnight at 4°C using a perm-and-block solution containing 0.5% Triton X-100, and 3% bovine serum albumin in PBS. Then mOLCs were incubated overnight at 4°C with primary rabbit anti-Influenza NP antibody (PA5-32242; Invitrogen) diluted 1:400 in the perm-and-block solution. After 3 washes in PBS, mOLCs were incubated overnight at 4°C in the perm-and-block solution with a secondary donkey anti-rabbit antibody labelled with Alexa Fluor 555 diluted 1:500 (A31572; Invitrogen). After 3 washes, mOLCs were incubated for 1 h at room temperature with Hoechst diluted 1:1000. After 3 washes, slices were mounted using Prolong antifade mounting medium. Immunofluorescence was imaged using Nikon Eclipse Ts2R Optical microscope.

### Metabolic activity of mOLCs

Evaluation of the metabolic activity of mOLC was carried out using a 3-(4,5-dimethylthiazol-2-yl)-2,5-diphenyltetrazolium bromide (MTT) assay as previously described (23). Briefly, mOLCs were collected and incubated for 4 h at 37°C with 200 µL of MTT (M2128; Sigma) at 5 mg/mL. Mitochondrial dehydrogenase enzymes reduce MTT into purple formazan crystals that can be quantified spectrophotometrically. Slices were transferred in a 96-well plate with 100 µL of DMSO and incubated for 10 min at 37°C for formazan crystals solubilization. Colored supernatant was then transferred and absorbance was measured at 540 nm.

### LDH release assay

To assess cellular cytotoxicity, LDH (lactate dehydrogenase) release was measured in mOLC culture medium using the LDH-Glo Cytotoxicity Assay (Promega). Media were collected and diluted 100 times in LDH storage buffer to be conserved at -20°C. The assay was performed according to the manufacturer’s instructions before measurement of luminescence with a Tristar 5 microplate reader (Berthold).

### Stereo-seq spatial transcriptomic analysis of mOLCs

Preparation of mOLCs and cryosections. STOmics Stereo-seq is a spatial transcriptomic platform that captures mRNA from tissue sections using stereo chips. This technology achieves nanoscale resolution with a spot diameter of 220 nm. To construct a comprehensive spatial transcriptome of infected mouse lungs, mOLCs were prepared and infected as described above. Two days post-infection mOLCs were collected and inactivated for 2 h with 4% PFA at room temperature, protected from light, followed by three washes in PBS, before being embedded in Tissue OCT matrix (KMA-0100-00A; Cell Path) and stored at -80°C until analysis. Tissue cryosections of 12 µm thickness were then performed using cryostat (CM1950; Leica) at -20°C and deposited on the Stereo-seq 1 cm x 1 cm transcriptomics chip for transcriptome analysis and dried for 5 min at 37°C according to manufacturer. Further steps were performed by ProfileXpert platform (Lyon, France). Three independent experiments were performed using the FF V1.2, FF V1.3 and FFPE V1.0 chemistries.

Stereo-seq library preparation and sequencing. The cryosections were fixed in pre-cooled methanol for 15 min at –20°C and stained with DAPI (0.02 µg/ml) for 5 min, followed by 3 washes in 0.1X SSC buffer. Imaging was performed with Zeiss Axioscan 7 at 10x objective for later nuclei segmentation. The chip was then incubated in decrosslinking buffer for 1 h at 70°C followed by another 15 min fixation step in pre-cooled methanol at –20°C. Subsequently, the tissue section was incubated in the permeabilization buffer for 15 min at 37°C. Captured RNAs from the tissue were then reverse transcribed for 5 h at 42°C. Next, the tissue was removed at 55°C for 10 min in Tissue Removal buffer and the cDNAs were released from the chip overnight at 55°C. After their size selection, amplification and purification, the concentration was quantified using the Qubit dsDNA HS assay kit. 20 ng of cDNA were used for library construction using the library preparation kit. For the FFPE chemistry only, an additional step of ribosomal depletion was performed using the SEQuoia Ribo Depletion Kit (Biorad) according to STOmics protocol (STUM-LP004.v.A-2).

Finally, the DNBs were sequenced on the MGI DNBSEQ G400 at the ProfileXpert genomic platform with 50 bp read1 and 100 bp read2 for FF chemistries and 25 bp read 1 and 60 bp read 2 for FFPE chemistry.

### Stereo-seq data analysis

Stereo-seq data processing. The fastq files were processed following the standard Stereo-seq pipeline SAW V8.2 provided by STOmics (https://github.com/STOmics/SAW). Reads containing coordinate identity (CID) sequences and molecular identifiers (MIDs) of good quality scores were aligned to the reference mouse genome GRCm38 mm10 customized with flu virus sequences using STAR. Only reads achieving a good mapping quality score were considered for gene annotation. Finally, the CID-containing expression count matrix was generated.

Bioinformatics analysis. Because one stereo-chip contained millions of spots (diameter: 220 nm), we merged adjacent 50×50 spots to one spot (bin50, 25 μm resolution) as a fundamental unit for downstream analysis, including unsupervised clustering. Using Scanpy package (V1.9.6) spots were quality-controlled by filtering low quality spots based on the number of detected genes (<2000), the total UMI counts (<5000) and the mitochondrial genes content (<10%) in each spot. We obtained 27 003 spots in total with a mean of 1000 genes and 2 350 UMI counts per spot. Spots expression matrix was then normalized using the log1p transformation.

Differential gene analysis. To identify changes in gene expression in infected tissue versus control mock tissue we performed a nonparametric Wilcoxon rank sum test using the scanpy.tl.rank_genes_groups() function from Scanpy (V1.9.6). P value adjustment was performed using benjamini-hochberg to identify significant DEGs with adjusted p value below 0.05. The lists of DEGs between the Mock and the infected slice for 3 distinct experiments were compared using a three-circle Venn diagram using BioVenn application (66).

### Statistical analyses

Data are presented as means ± SEM. At least 3 independent experiments were analyzed and the number of replicates is specified in the figure legends. All statistical analyses were carried out using GraphPad Prism 10. Comparison of two groups was performed using Student’s t-test. For comparisons involving multiple groups, a one-way ANOVA or Wilcoxon test (for normalized data) was used, and p-values were adjusted for multiple comparisons using post-hoc tests or analysis of stacks of p-values. Outliers were identified and excluded using the ROUT method (Q = 0.05) in GraphPad Prism. p*<0.05, **p<0.005, ***p<0.0005, ****p<0.0001. ns: not significant; nd: not determined (no variance).

## Acknowledgments

We thank Déborah Brea-Diakité (CEPR, Tours) for sharing H3N2 Influenza A/Scotland/20/74 strain. We thank Emma Teissedre for technical assistance during her Master1 internship. We acknowledge the contribution of SFR Santé Lyon-Est (UAR3453 CNRS, US7 Inserm, UCBL) facility: CIQLE (a LyMIC member), especially Bruno Chapuis for his help in image acquisition of Stereo-Seq spatial transcriptomics slides. We acknowledge the contribution of SFR Biosciences (Université Claude Bernard Lyon 1, CNRS UAR3444, Inserm US8, ENS de Lyon) facilities of AniRA-PBES and AniRA-ImmOs metabolic phenotyping, and especially thank Romain Barnault, Laurence Canaple and Sophia Djebali for their help. This research was funded by an intramural CIRI grant to C.M. and L.P.-C., the Agence National de la Recherche / Agence Innovation Défense / Direction Générale de l’Armement (ViroMetaBlock project; ANR-22-ASTR-0021 to L.P.-C., C.M., V.L., and M.S.-T., ANR-22-CE15-0001 to PO.V. and ANR-24-CE15-3934-03 to L.P.-C.). FJ was funded by a Ph.D fellowship from the Agence Nationale de la Recherche ANR-22-CE15-0001-01 and a post-doctoral fellowship from the Fondation CNRS - Vidalain. LC was funded for her Ph.D by INSERM, the Direction Générale de l’Armement (DGA), and the Agence Innovation Défense (AID).

## Author Contributions

F.J, A.A-G, M.V., D.D., L.C., E.O., A.C., C.J, G.P., S.C., G.M., C.M performed experiments and analyzed data. M.L., T.D. and G.M. analyzed data. J.L. supervised spatial transcriptomic platform. L.P–C. designed the study and conducted the research with the help of C.M., P-O.V., and O.D. F.J., A.A-G and L.P-C. wrote the first draft of the manuscript. P-O.V., O.D. C.M., M.S-T., G.M. and V.L contributed to the building of the project and revised the manuscript.

## Competing Interest Statement

A patent application related to this work has been submitted.

## Notes

### Competing Interest Statement

The authors have declared no competing interest.

### Summary of Updates

In the revised version of this paper, we have included novel data of spatial RNAseq on infected mOLCs with advanced analysis. We have strengthened the link between intracellular NAD⁺ content, protein MARylation and PARP activity. Furthermore, additional experiments characterize the metabolic consequences of NAD depletion (new Fig. 2) and establish an association between NAD availability, protein MARylation and viral replication (new Fig. S12). In addition, we have better characterized the mode of action of the antiviral molecules targeting NAD biosynthesis (FK-866) and NAD consumption by PARPs (OUL232). Importantly, we provide novel ex vivo results with combined OUL232 and FK-866 treatments, that strongly reduced the MARylation activity in cells (see new Fig S12), and IAV production in lungs by 2 log10 (see new Fig. 5D)

