## Supplementary material for "Combination of in vitro and ex vivo infection models highlights the role of NAD-dependent ADP-ribosylation in Influenza A virus infection": SI appendix

[*]Dr. Laure Perrin-Cocon, CIRI, VIRIMI team, 21 av. Tony Garnier, 69365 Lyon, France.

**This PDF file includes:**

Figures S1 to S14

Legend for Dataset S1

SI References

**Other supporting materials for this manuscript include the following:**

Dataset S1

### Figures

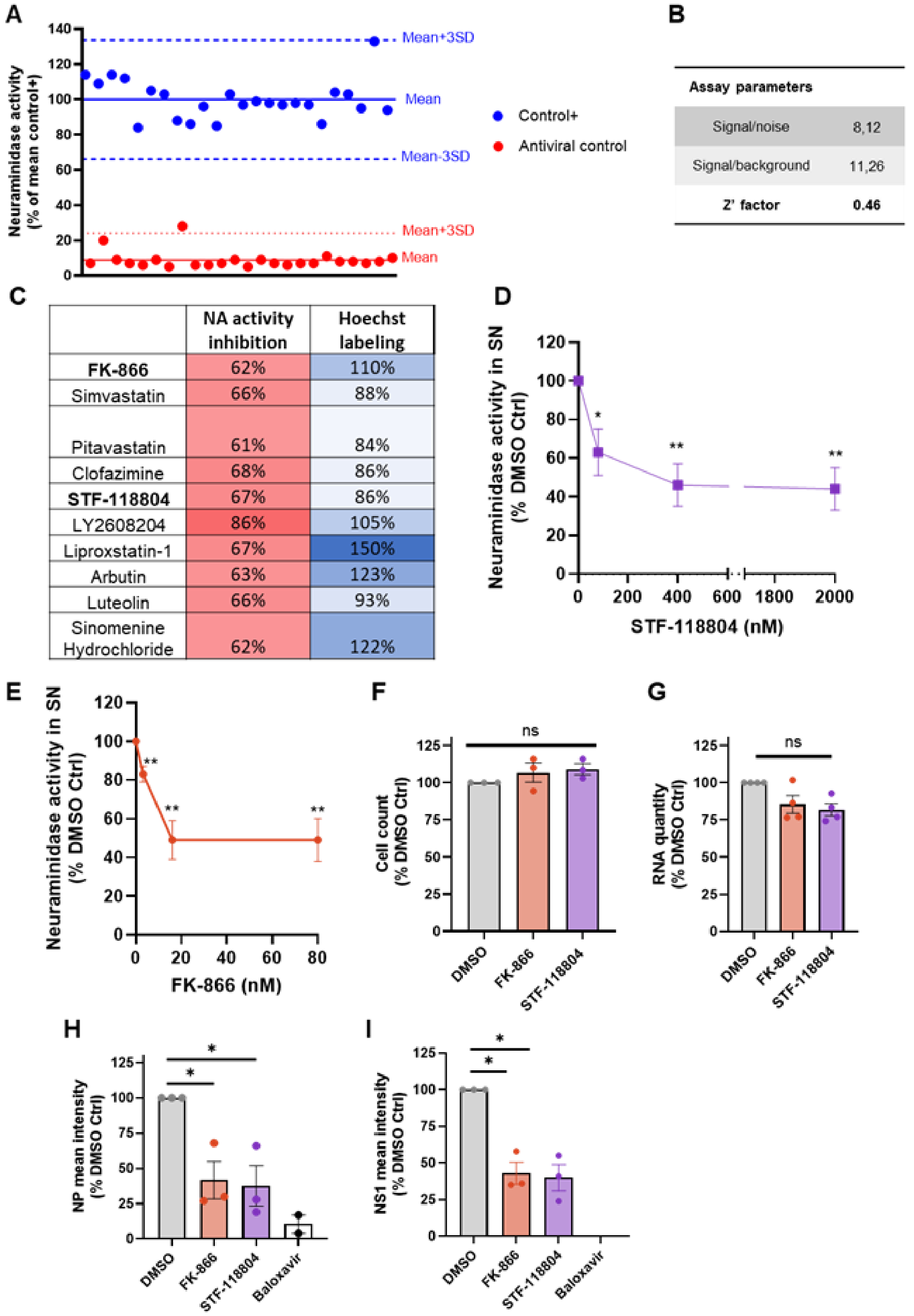


**Fig. S1.** NAMPTi restrain IAV infection. (A-B) Quality controls of the screening. A549 cells were seeded and pretreated for 24 h with 20 nM Baloxavir (antiviral control) or DMSO solvent (positive control). Influenza A/H3N2/Scotland/20/1974 virus was introduced at MOI 0.5 without medium change, and 48 hpi viral production was determined by neuraminidase activity assay in supernatants. Positive and antiviral controls were placed in quadruplicates on each plate. (A) Neuraminidase activity was measured in cells supernatant and presented as a percentage of the mean value of all the positive controls. Distribution of neuraminidase activity of every well of infected positive and antiviral controls measured across the 6 plates of the screening. Means and means ± 3SD representing the "hit threshold" as described by Zhang et al. are shown(1). (B) Quality of this screening is also described by the following assay parameters:

Signal/noise= $\frac{mean\ (positive\ controls) - mean\ (antiviral\ controls)}{standard\ deviation\ (positive\ controls)}$

Signal/background: $\frac{mean\ (positive\ controls)}{mean\ (antiviral\ controls)}$

Z' factor: $1 - \frac{3SD\ (positive\ controls) + 3SD\ (antiviral\ controls)}{|mean\ (positive\ controls) - mean\ (antiviral\ controls)|}$

(C) Results of the screening. List of the drugs reducing NA activity by at least 60% that did not reduce the cell count by more than 20% compared to DMSO.
(D-E) Inhibitory effect of increasing doses of NAMPT inhibitors on NA secretion. Experiments were carried out as described (A) with 80, 400 and 2 000 nM of STF-118804 (D) or 3.2, 16 or 80 nM of FK-866 (E). Data correspond to means ± SEM of at least 3 experiments in triplicates.
(G) Total RNA quantity extracted at 48 hpi from infected cells pretreated with 80 nM of FK-866 2 µM of STF-118804 or DMSO. Data shown are means ± SEM of 4 experiments.
(H-I) Expression of viral nucleoprotein NP (D) or non-structural protein NS1 (E) measured after immunostaining at 4 h post-infection. Cells were pretreated as in G. Data are the mean fluorescent intensities quantified on 3 views/well in triplicates. Means ± SEM of 3 independent experiments is shown.

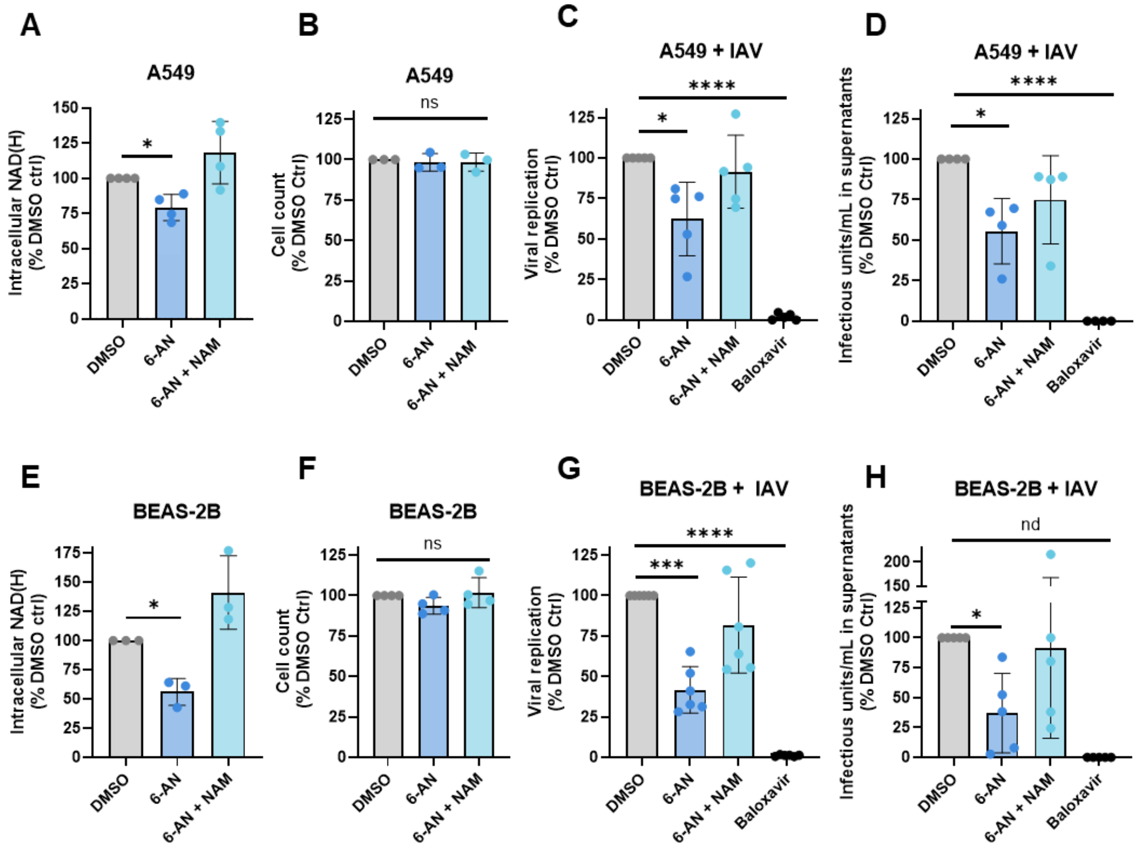


**Fig. S2.** Interfering with NAD pathway using 6-AN restrains IAV replication and production of infectious particles in vitro. A549 cells (A-D) or BEAS-2B cells (E-H) were treated with 6-AN (100 µM) supplemented or not with nicotinamide (NAM; 500 µM) for 24 h. Intracellular NAD(H) was quantified (A, E) and cell count determined by Hoechst nuclei staining (B, F). Cells were infected with IAV H3N2 at MOI 1 for 1.5 h before removal of viral inoculum and treatment with Baloxavir (20 nM) or 6-AN (100 µM) with or without nicotinamide (500 µM). Supernatants and cells were collected 24 h post-infection. Expression level of M viral segment in cell lysates was quantified by RT-qPCR and normalized with mRNA quantities of Rplp0 cellular gene (C, G). Infectious viral particles concentration in cells supernatants was determined by TCID50 assay on MDCK cells (D, H). Data shown are means ± SEM of independent experiments. ns: not significant; nd: not determined (no variance).

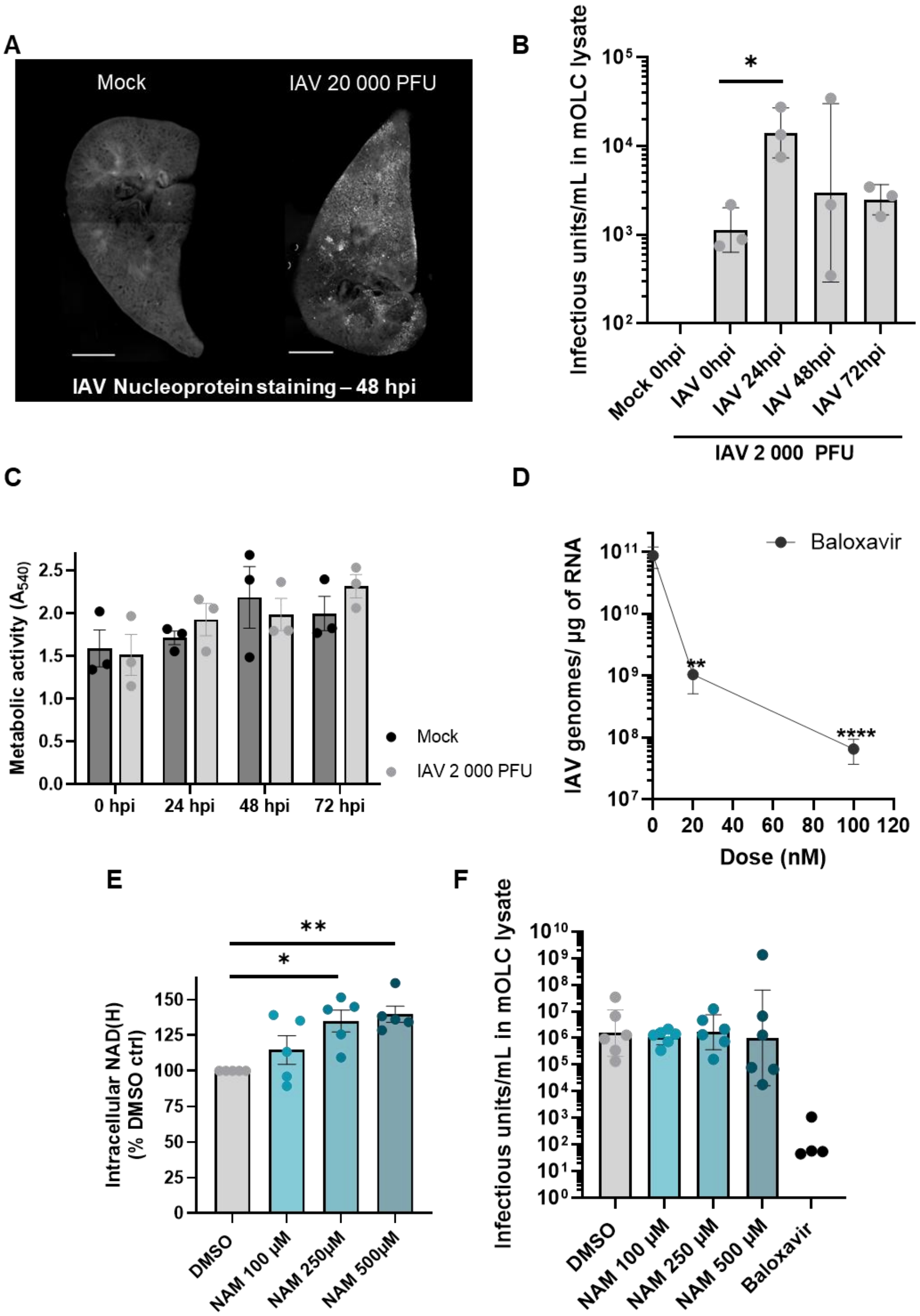


**Fig. S3.** mOLC model supports IAV replication and viral production over time
mOLCs were infected or not (mock) with 2 000 or 20 000 PFU of IAV H3N2, as indicated.
(A) Immunofluorescent staining of viral nucleoprotein at 48hpi with 20 000 PFU. mOLCs were observed using x4 objective (100 ms of exposure) and the picture of the entire slice was reconstituted using the image stitching function of Image J. Scale bar = 1 mm.
(B) Concentration of viral infectious particles in lysates of mOLCs infected with 2 000 PFU was determined just after (0), 24, 48 or 72 h post infection by $TCID_{50}$ assay on MDCK cells. Data from 3 independent animals are shown with geometrical means ± SD.

(C) Metabolic activity was measured just after (0), 24, 48 or 72 h post infection by MTT assay. Data from 3 independent animals with means ± SEM.
(D) The amount of viral M segment in mOLCs lysates was determined by RT-qPCR at 48 h after infection with 20 000 PFU of IAV and treatment with increasing doses of Baloxavir (0, 20 and 100 nM). Data from 4 independent animals with geometrical means ± SD.
(E-F) mOLCs were infected with 2 000 PFU of IAV before being treated with nicotinamide (NAM 100, 250 or 500 µM). Treatment of mOLCs as well as medium were renewed at 24 hpi. Slices were collected at 48 hpi. (E) Three mOLCs from different animals were pooled for lysis and NAD(H) assay. Individual data are presented with means ± SEM. (F) Concentration of viral infectious particles in individual mOLC lysate was determined by $TCID_{50}$ assay on MDCK cells. Geometrical means ± SD of 4-6 independent animals is shown.

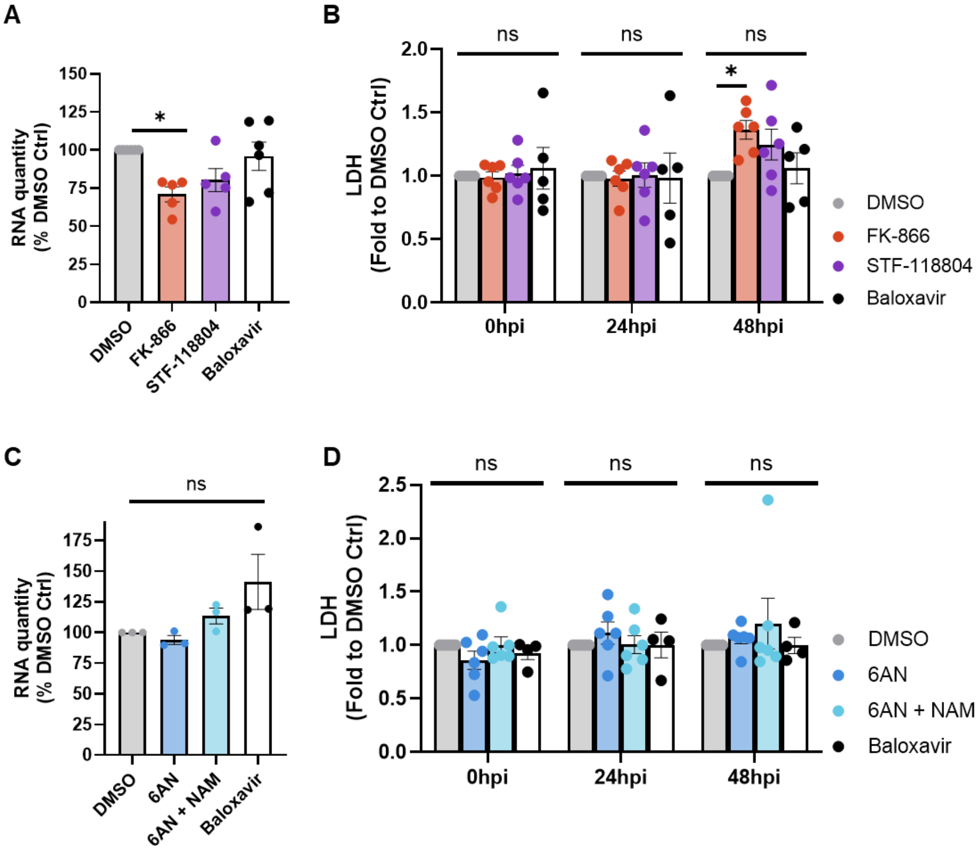


**Fig. S4.** mOLC viability upon exposure to NAD regulators.
mOLCs were infected with 2 000 PFU of IAV H3N2 before being treated with NAMPTi (20 µM for FK-866 and STF-118804), 6-AN (100 µM), 6-AN with nicotinamide (NAM; 500 µM) or IAV inhibitor Baloxavir (20 nM). mOLC treatment and medium were renewed at 24 hpi.
(A, C) RNA quantity extracted from 3 mOLCs from different animals at 48 hpi. Data are means of minimum 3 experiments.
(B, D) Lactate dehydrogenase released in the medium by 3 mOLCs from different animals at 0, 24 and 48 hpi. Data represented are the means of minimum 3 independent experiments.

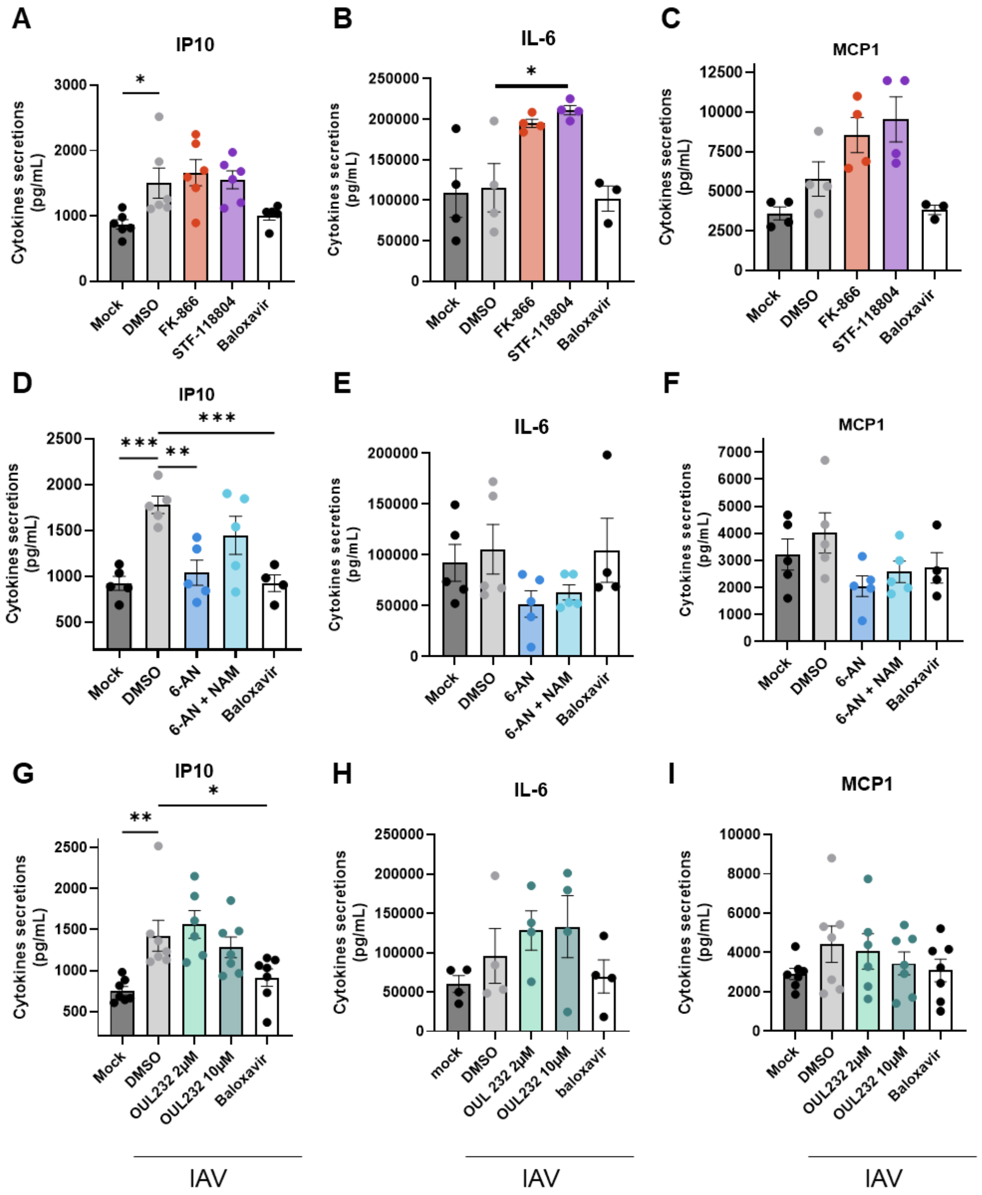


**Fig. S5.** Modulation of cytokines secretion upon infection and treatments of mOLCs. mOLCs were infected with 2 000 PFU of IAV before being treated with NAMPTi (20 µM for FK-866 and STF-118804) (A-C), 6-AN (100 µM), 6-AN with nicotinamide (NAM; 500 µM) (D-F), OUL232 (2 or 10 µM) (G-I) or IAV inhibitor Baloxavir (20 nM) (A-I). mOLC treatment and medium were renewed at 24 hpi, before collection at 48 hpi. Cytokine concentrations were determined in mOLCs subnatants, using CBA Flex multiplex assay for IP-10 and MCP1 and MSD U-PLEX for IL-6.

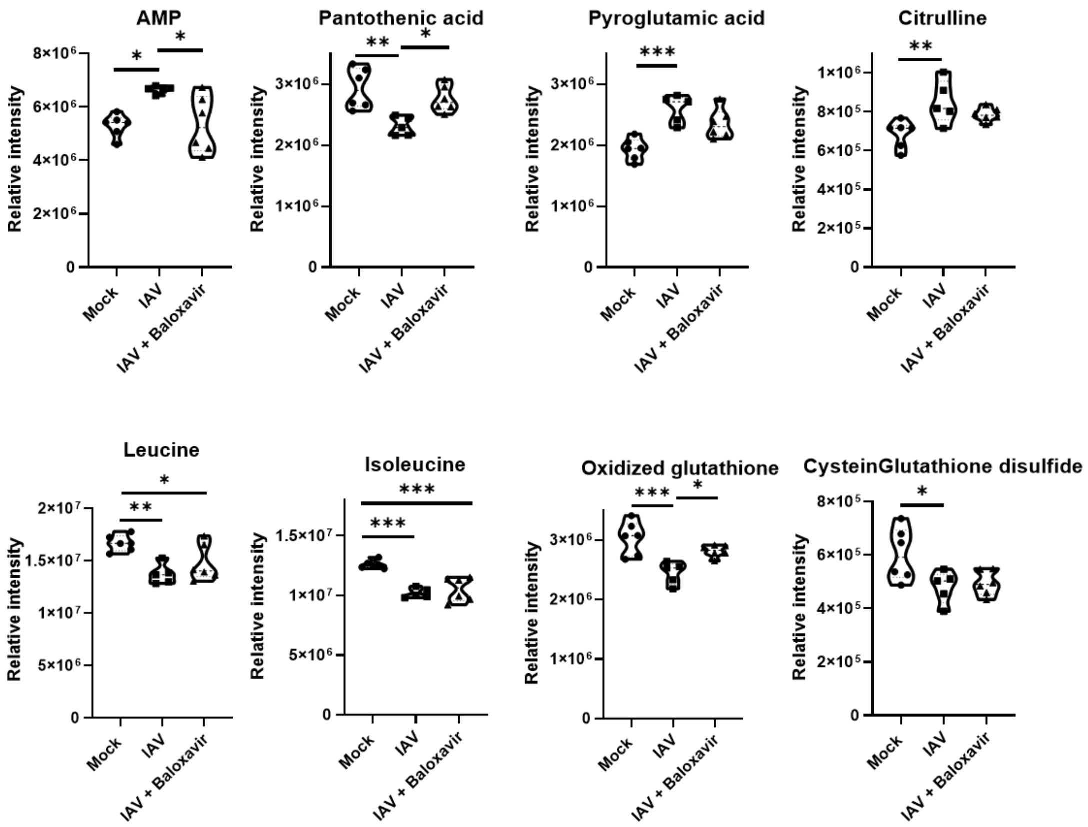


**Fig. S6.** IAV infection induces metabolic changes in mOLCs, some are reverted by treatment with IAV inhibitor Baloxavir.

Metabolic profile of mOLC 48 hpi, expressed as relative peak intensities in mOLCs not infected (mock), infected with IAV 20 000 PFU (IAV) and infected and treated with 100 nM of Baloxavir (IAV + Baloxavir). Two distinct pools of 5 mOLCs from independent mice were measured in triplicate for metabolite analysis by LC-MS/MS semi-quantitative analysis. Relative intensities of metabolites significantly modulated by IAV infection compared to mock in mOLCs ($p<0.05$).

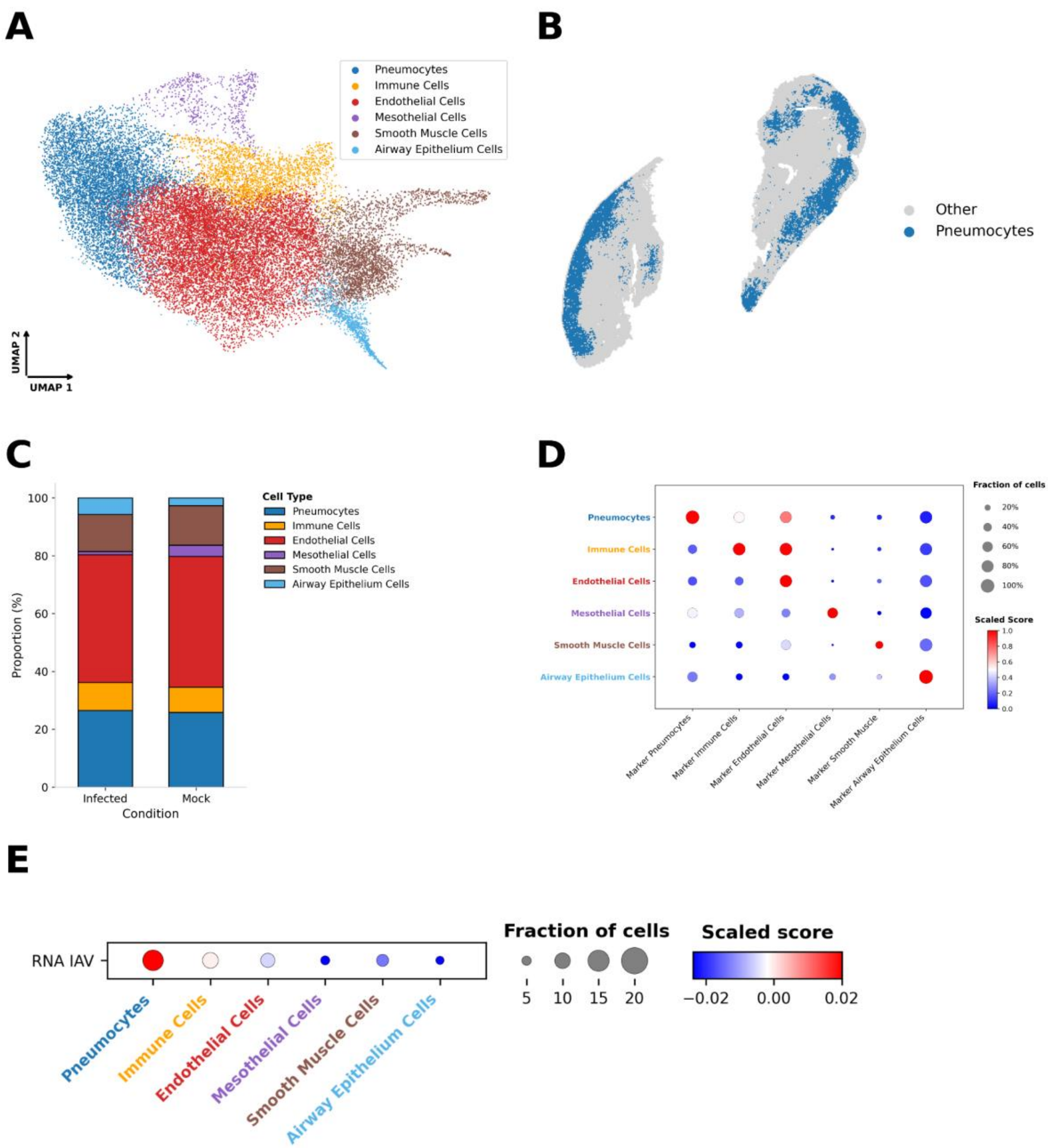


**Fig. S7.** Identification and annotation of lung cell types and viral RNA expression. (A) UMAP embedding showing the distribution of the identified cell clusters. (B) Spatial distribution of the pneumocytes cluster. (C) Proportions of the different cell clusters in the mock and infected slice. (D) Dot plot validating the expression profiles of canonical marker genes used to annotate the major lung cell types. (E) Dot plot displaying the expression of viral RNAs across the different cell clusters. To ensure that clustering was driven solely by intrinsic cell type identity and not biased by infection-induced transcriptional changes, unsupervised clustering was first performed exclusively on the mock slice using a nearest-neighbor graph derived from the principal component analysis (PCA) of highly variable genes (HVGs). This healthy reference clustering was then projected onto the two other slices using an ingest procedure. The resulting clusters were subsequently refined and merged based on correlation matrices and the preservation of marker gene signatures to define the final major lung cell populations.

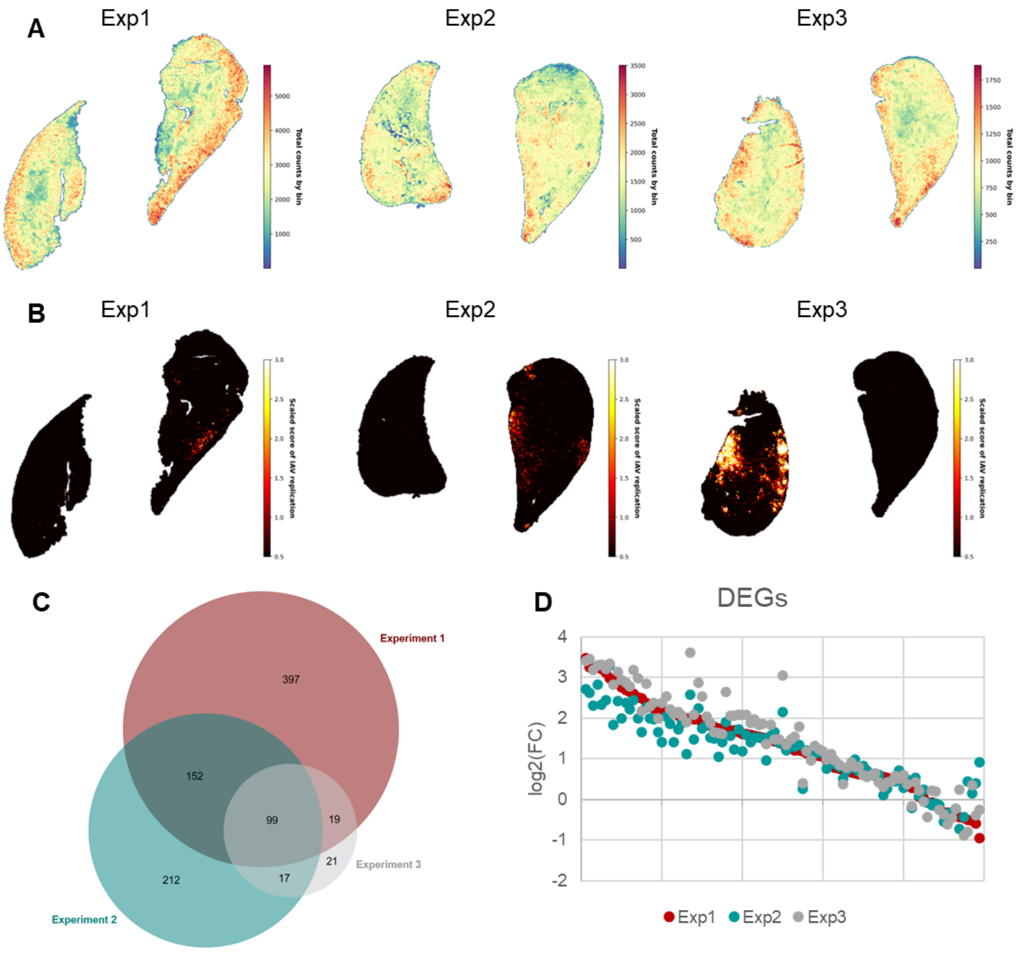


E

| GO-Term BP direct | Number of genes | Fold Enrichment | FDR |
|---|---|---|---|
| **Innate immune response** | 25 | 24,67 | $3,31\ 10^{-26}$ |
| **Defense response to virus** | 20 | 48,84 | $9,23\ 10^{-26}$ |
| **Immune system process** | 22 | 15,07 | $2,52\ 10^{-18}$ |
| **Cellular response to interferon-beta** | 11 | 95,72 | $8,35\ 10^{-16}$ |
| **Response to bacterium** | 14 | 30,24 | $1,91\ 10^{-14}$ |
| **Response to virus** | 11 | 59,10 | $8,52\ 10^{-14}$ |
| **Activation of innate immune response** | 9 | 104,92 | $5,89\ 10^{-13}$ |
| **Antiviral innate immune response** | 9 | 63,92 | $3,27\ 10^{-11}$ |
| **Negative regulation of viral genome replication** | 7 | 92,02 | $3,12\ 10^{-09}$ |
| **Response to other organism** | 5 | 154,47 | $6,54\ 10^{-07}$ |

**Fig. S8.** Analysis of three distinct experiments by Stereo-Seq spatial transcriptomic study. (A) Heatmap of the total number of reads. (B) Heatmap of all the viral transcripts detected. (C-D) Differentially expressed genes (DEGs) between infected and mock mOLCs across three independent Stereo-Seq experiments. BioVenn **(2)** was used to analyze the list of genes differentially expressed (p<0,05) in the 3 distinct experiments (C). Comparison of the log2(fold change; FC) of the 99 DEGs conserved across the 3 experiments (D). (E) Analysis of the functional annotation of the 99 conserved DEGs between infected and mock mOLCs: TOP 10 significantly enriched biological processes (GO terms-BP direct) in the murine DEGs.

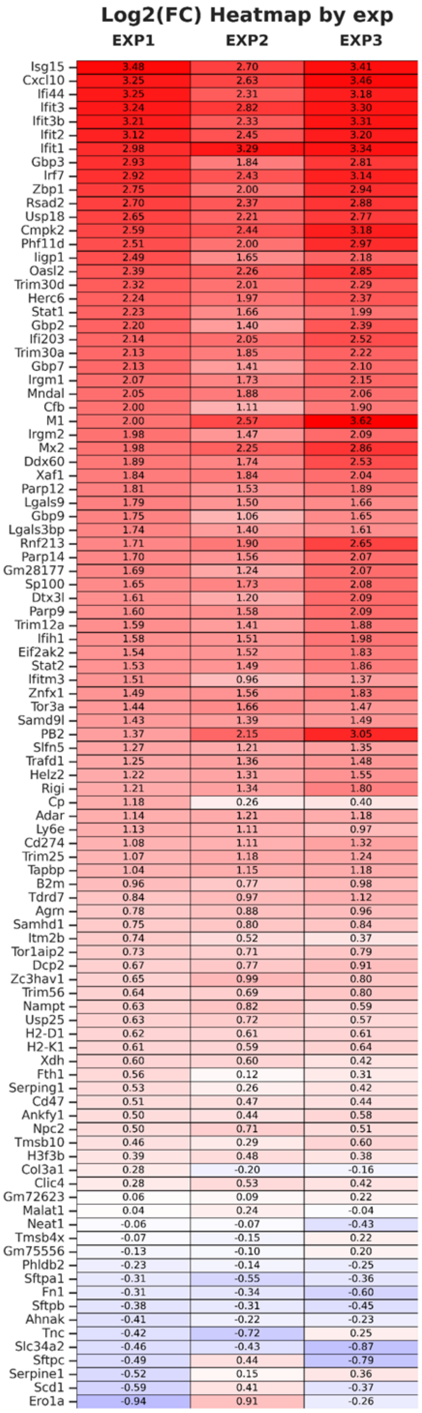


**Fig. S9.** Heatmap of significantly differentially expressed genes conserved across 3 independent experiments. For the 99 genes present at the intersection of the 3 experiments (cf. Venn Diagram, Figure S8), $log_2$(FC) between the infected and the mock slice is indicated for each experiment (EXP).

A

| NAD⁺ Synthesis Enzyme | Normalized counts/slice Mock | Normalized counts/slice Infected |
|---|---|---|
| Nampt | 3096 | 4752 |
| Nmnat1 | 171 | 201 |
| Nmnat2 | 327 | 325 |
| Nmnat3 | 321 | 316 |
| Naprt | 182 | 178 |
| Nadsyn1 | 125 | 135 |
| Qprt | 10 | 10 |

B

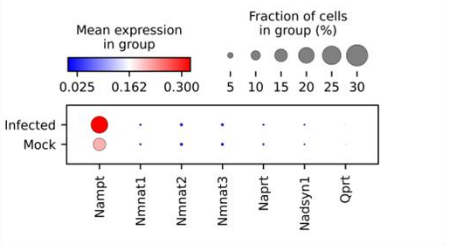


C

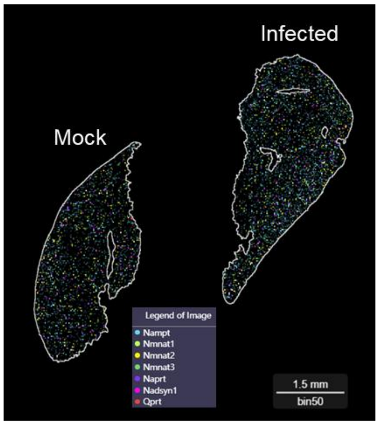


D

| NAD⁺ Consuming Enzyme | Normalized counts/slice Mock | Normalized counts/slice Infected |
|---|---|---|
| Sirt1 | 648 | 597 |
| Sirt2 | 1429 | 1546 |
| Sirt7 | 1104 | 1181 |
| Cd38 | 3595 | 3413 |
| Sarm1 | 243 | 220 |
| Parp9 | 934 | 2918 |
| Parp12 | 1035 | 3596 |
| Parp14 | 1628 | 5326 |

E

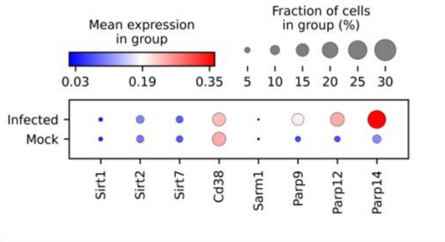


F

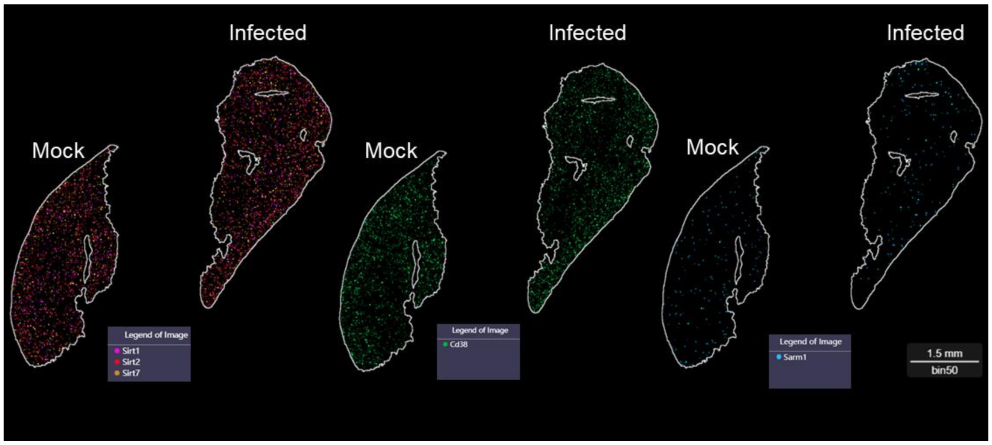


**Fig. S10.** Stereo-Seq analysis of genes involved in NAD⁺ biosynthesis and consumption. (A, D) Normalized counts of transcripts detected by Stereo-Seq for some enzymes involved in NAD⁺ biosynthesis (A) and NAD⁺ consumption (D).
(B, E) Dot plots presenting the intensity of expression and the fraction of positive cells for the indicated genes.

(C, F) Spatial distribution of transcripts of $NAD^+$ biosynthesis (C) and $NAD^+$ consuming (F) enzymes in mock or infected mOLC cryosections.

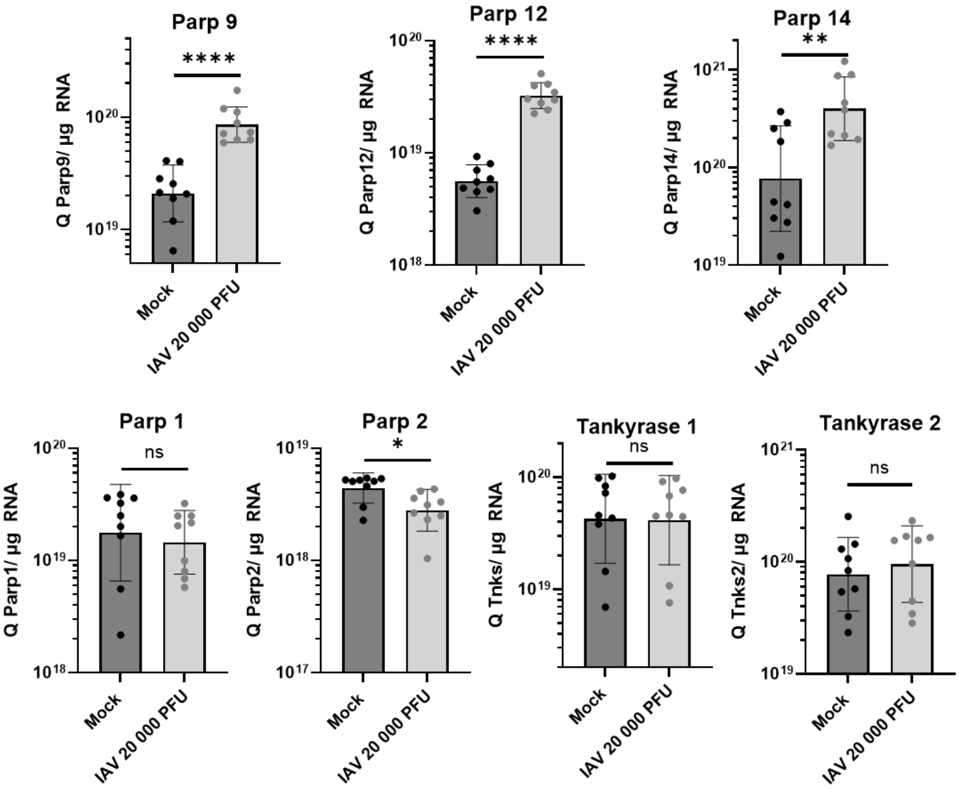


**Fig. S11.** Gene expression levels of PARPs monitored by RT-qPCR in IAV infected versus Mock mOLCs. mOLCs were infected or not (mock) with 20 000 PFU of IAV H3N2 for 48h. Each mOLC was lysed and RNA extracted for quantification by RT-qPCR. Gene expression was quantified using the $2^{-\Delta\Delta Ct}$ method and normalized to the housekeeping gene (Rplp0).

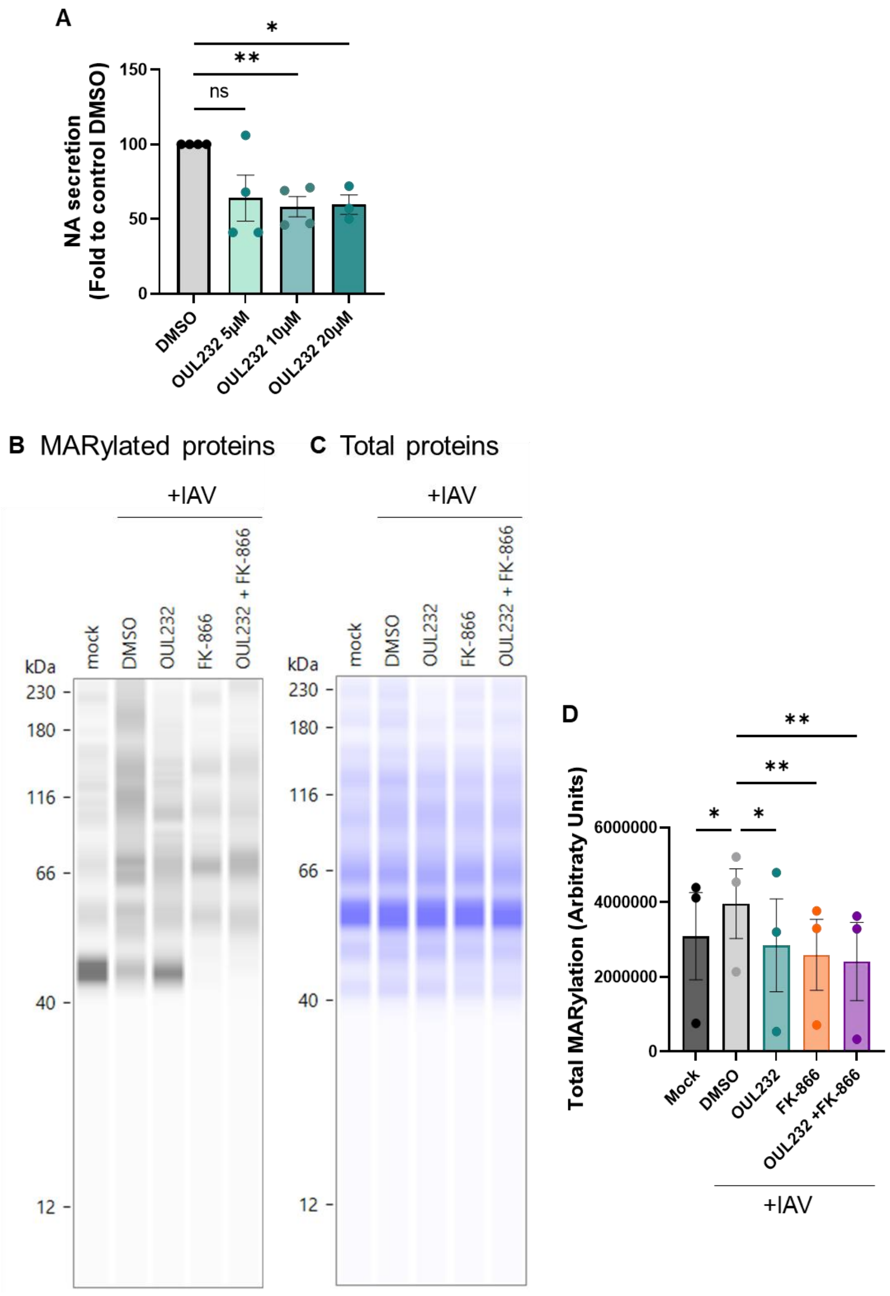


**Fig. S12.** OUL232 inhibitor restrains IAV viral infection and MARylating activity *in vitro*.

A549 cells were pretreated for 24 h with OUL232 at the indicated concentration (A) or 10 µM (B-D), 80 nM FK-866 with or without OUL232, or with control DMSO solvent, before being infected with IAV at MOI 0.1 for 24 h. (A) Viral production was determined by neuraminidase (NA) activity assay in supernatants. (B-D) Cells were treated by MG132 2 hpi to prevent degradation of MARylated proteins by the proteasome, and harvested at 24 hpi. MARylated proteins were detected in cell lysates by simple Western (B) and total proteins were stained on the same capillary after Re-Plex and used for normalization (C). The total amount of MARylated proteins, normalized to total protein abundance, was quantified using Compass Software by integrating the area under the curve of the spectra (D). Statistical analysis was performed using one-way ANOVA for paired samples.

**A**

| IC50 (µM) | PARylating PARPs | | | | MARylating PARPs | | | | |
|---|---|---|---|---|---|---|---|---|---|
| **Inhibitor** | **PARP1** | **PARP2** | **TNKS1** | **TNKS2** | **PARP7** | **PARP10** | **PARP11** | **PARP12** | **PARP14** |
| OUL-232 | 15[a] | 10[a] | 5.4[a] | 10[a] | 0.083[a] | 7.8 nM[a] | 0.24[a] | 0.16[a] | 0.3[a] |
| OUL-35 | >20[b] | | | | 5.5[b] | 0.33[a]; 3.3[b] | | 6.8[b] | |
| PARP10/15-IN1 | >100[c] | >100[c] | >100[c] | >100[c] | | 0.16[c] | | >10[c] | >10[c] |
| H-10 | 11.68[g] | | 9.04[g] | | | | | | 0.49[d,g] |
| Olaparib | <1 nM[b]; 1-19 nM[e]; 1 nM[f] | 0.3 nM[e]; 0.001-0.25[f] | 1.23-3.5[f] | 1.7-2.34[f] | 0.48[b] | 8.2[b]; 0.86[e]; 1.25-9.3[f] | | 7.5[b]; 3.8[e]; 10.05[f] | >100[e] |
| Nesuparib (JPI-547) | 0.002[h] | | 0.005[h] | 0.001[h] | | | | | |

less than 0.01 µM
from 0.01 to 1 µM
from 1 to 20 µM
more than 20 µM

a Murthy S et al., J Med Chem 2023
b Lu AZ et al., Biochem Pharmacol 2019
c Nizi MG et al., Eur J Med Chem 2022
d Wu S et al., J Med Chem 2025
e Wang L-M et al., EMBO Mol Med 2023
f Antolin AA et al., Sci Rep 2020
g Peng B et al., Angew Chem Int Ed 2017
h Yu M et al., J Med Chem 2022

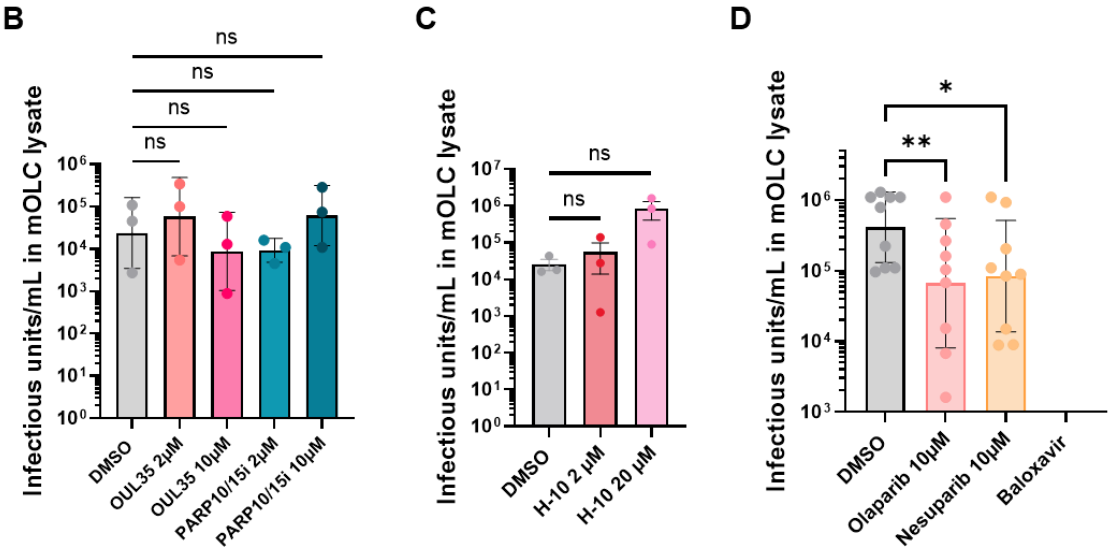


**Fig. S13.** Effect of several PARP inhibitors on IAV infection *ex vivo*.

(A) Table of published IC50 values for PARP inhibitors used in this publication (3–10). (B-D) mOLCs were infected with 2 000 PFU of IAV H3N2 before being treated with the indicated concentration of OUL35, PARP10/15i, H-10, Olaparib, Nesuparib or 100 nM Baloxavir. Treatments and media were renewed at 24 hpi and slices were collected at 48 hpi. The production of infectious viral particles in mOLC lysates was determined by $TCID_{50}$ assays on MDCK cells. (D) A549 cells were pretreated for 24 h with PARP inhibitors at the indicated concentration or with control DMSO solvent, before being infected with IAV at MOI 0.1 for 24 h. Viral production was determined by neuraminidase (NA) activity assay in supernatants.

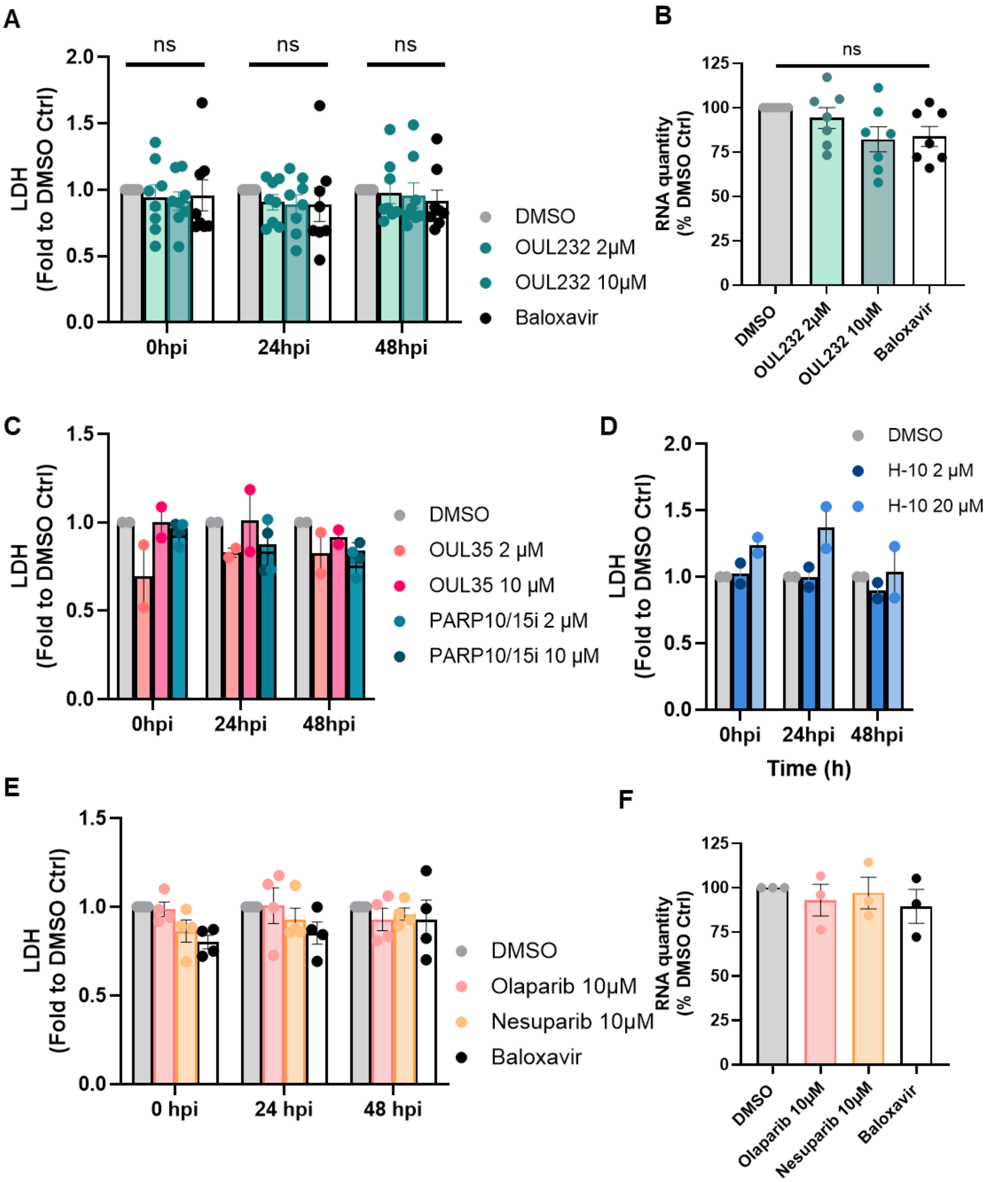


**Fig. S14.** Viability of mOLCs upon PARP inhibitor treatment.
mOLCs were infected with 2 000 or 20000 PFU of IAV H3N2 before being treated with OUL232, OUL35, PARP10/15i, H-10, Olaparib or Nesuparib at the indicated concentration. Treatments and media were renewed at 24 hpi.
(A,C,D,E) Lactate dehydrogenase (LDH) mean release in the medium by 3 mOLCs from different animals at 0, 24 and 48 hpi. Mean data from independent experiments and means ± SEM are presented.
(B,F) RNA quantity extracted from mOLCs from different animals at 48 hpi. Individual data and means ± SEM.

**Dataset S1 (separate file).** Liste of genes significantly modulated between the infected and the mock slice ($p<0,05$) for each experiment (Exp), with log2(fold change; FC).
